# Imprinted *Grb10*, encoding growth factor receptor bound protein 10, regulates fetal growth independently of the insulin-like growth factor type 1 receptor (*Igf1r*) and insulin receptor (*Insr*) genes

**DOI:** 10.1101/2024.01.24.576998

**Authors:** Kim Moorwood, Florentia M. Smith, Alastair S. Garfield, Andrew Ward

**Affiliations:** University of Bath, Department of Life Sciences, Building 4 South, Claverton Down, Bath, BA2 7AY, United Kingdom

## Abstract

**Background:** Optimal size at birth dictates perinatal survival and long-term risk of developing common disorders such as obesity, type 2 diabetes and cardiovascular disease. The imprinted *Grb10* gene encodes a signalling adaptor protein capable of inhibiting receptor tyrosine kinases, including the insulin receptor (Insr) and insulin-like growth factor type 1 receptor (Igf1r). *Grb10* restricts fetal growth such that *Grb10* knockout (KO) mice are at birth some 25-35% larger than wild type. Using a mouse genetic approach, we test the widely held assumption that Grb10 influences growth through interaction with Igf1r, which has a highly conserved growth promoting role.

**Results:** Should Grb10 interact with Igf1r to regulate growth *Grb10*:*Igf1r* double mutant mice should be indistinguishable from *Igf1r* KO single mutants, which are around half normal size at birth. Instead, *Grb10*:*Igf1r* double mutants were intermediate in size between *Grb10* KO and *Igf1r* KO single mutants, indicating additive effects of the two signalling proteins having opposite actions in separate pathways. Some organs examined followed a similar pattern, though *Grb10* KO neonates exhibited sparing of the brain and kidneys, whereas the influence of *Igf1r* extended to all organs. An interaction between Grb10 and Insr was similarly investigated. While there was no general evidence for a major interaction for fetal growth regulation, the liver was an exception. The liver in *Grb10* KO mutants was disproportionately overgrown with evidence of excess lipid storage in hepatocytes, whereas *Grb10*:*Insr* double mutants were indistinguishable from *Insr* single mutants or wild types.

**Conclusions:** Grb10 acts largely independently of Igf1r or Insr to control fetal growth and has a more variable influence on individual organs. Only the disproportionate overgrowth and excess lipid storage seen in the *Grb10* KO neonatal liver can be explained through an interaction between Grb10 and the Insr. Our findings are important for understanding how positive and negative influences on fetal growth dictate size and tissue proportions at birth.

## Introduction

Mammalian fetal growth is a highly regulated process influenced positively and negatively by genetic and environmental factors, including maternal nutrient supply. Attaining an appropriate size is strongly correlated with infant survival [1] and minimises the risk in later life of common disorders including obesity, diabetes and cardiovascular disease (see [2, 3]). The insulin/insulin-like growth factor (Ins/IGF) signalling pathway is conserved, most likely throughout animal species, to regulate growth and energy homeostasis, as well as being a major determinant of longevity [4, 5]. Involvement of TOR (mTOR in mammals) is similarly broadly conserved and links nutrient sensing and protein translation control with the same processes [5]. The invertebrate pathway involves a single Ins/Igf receptor that mediates all of these functions. In mammals the regulation of energy metabolism is a separate function of insulin acting through the insulin receptor (Insr), while the Igf1r is the primary mediator of fetal growth, stimulated by the Igf1 and Igf2 ligands (Figure 1A). This was established through a series of elegant mouse genetic experiments that also linked fetal growth regulation with genomic imprinting [6].

**Figure 1.**
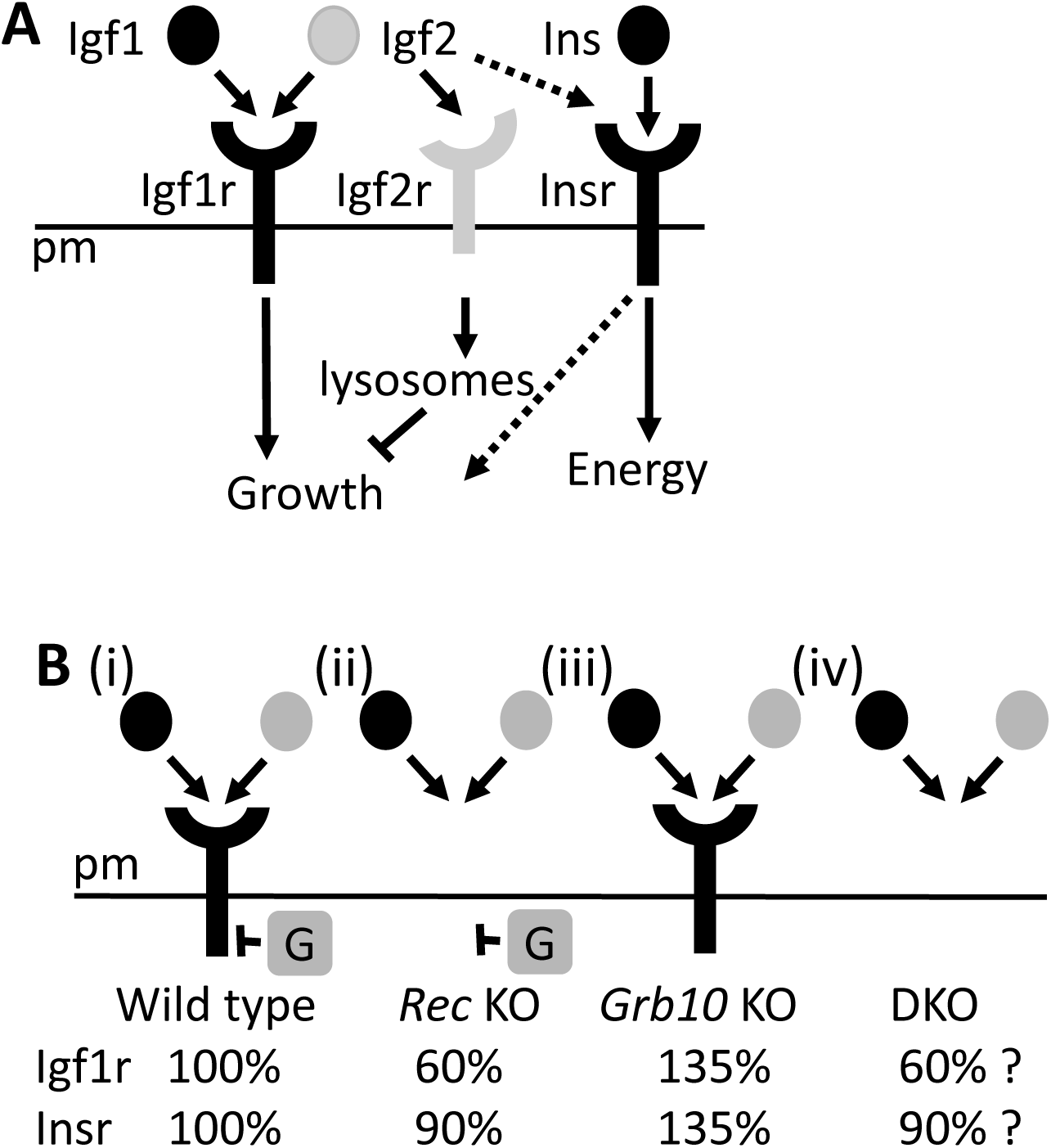
Signalling interactions within the insulin/insulin-like growth factor pathway inferred from biochemical and mouse genetic studies. A) The Igf1 and Igf2 ligands bind and activate Igf1r to promote fetal growth, whereas insulin (Ins) activates the Insr predominantly to regulate energy homeostasis receptors. In the placenta, Igf2 also binds Insr, though with lower affinity than it does the structurally related Igf1r, to promote fetal growth. Igf2 is also bound by Igf2r and thereby targeted for lysosomal degradation, such that Igf2r has an inhibitory action on fetal growth through sequestration of Igf2. Products of imprinted genes, paternally expressed *Igf2* and maternally expressed *Igf2r*, are shaded (grey). B) Fetal growth outcomes expressed as mass at birth in mice of genotypes relevant to this study. Knockouts of either the *Igf1r* (i) or *Insr* (ii) previously shown to be growth restricted to 60% [15] and 90% [17] the size of wild type animals, respectively, while *Grb10* KO (iii) pups are enlarged at 135% [36–38]. If Grb10 should act predominantly on either receptor to inhibit growth then double knockout (DKO) mice, generated in the present study, should be indistinguishable from the respective receptor single KO pups (iv).

Genomic imprinting is a form of epigenetic gene regulation that relies on differential methylation of genes, established in the male and female germlines, that restricts expression to only one of the two parental alleles.The mouse genome contains around 150 imprinted genes, with just over half expressed predominantly from the paternally inherited allele and the rest expressed from the maternally inherited allele [7, 8]. Imprinted genes are diverse in their functions and the products they encode, but notable among them are genes encoding signaling proteins that regulate growth of the fetus, placenta or both. These genes tend to fit with the most widely accepted hypothesis for the evolution of genomic imprinting in mammals, which posits a conflict between parental alleles in offspring that can influence nutrient acquisition from the mother [9, 10]. Noting that a female may have multiple mates, it is in the father’s interest to maximise fitness of his offspring in an opportunistic manner, whereas the mother favours a more even distribution of resources to offspring throughout her reproductive span. These pressures have resulted in the expression in developing offspring of growth-promoting genes from paternally inherited alleles, such as *Igf2* and *Dlk1*, and growth restricting genes from maternally inherited alleles, such as *Cdkn1c*, *Grb10*, *Igf2r* and *Phlda2* [11, 12].

The first two imprinted genes to be identified encoded components of the insulin-like growth factor (IGF) signalling pathway (Figure 1A) [6]. Mouse knockout (KO) experiments were crucially important in identifying fetal growth regulatory functions of genes encoding the Igf1 and Igf2 ligands, and also the IGF type 1 (Igf1r) and type 2 (Igf2r) receptors. Separate knockouts of three different pathway components resulted in offspring that were markedly small at birth relative to wild type littermates, with *Igf1* KO pups approximately 60%, *Igf2* KO 60% and *Igf1r* KO 45% of normal size by mass [13–16]. Additionally, loss of insulin receptors had a relatively modest impact on growth, with *Insr* KO pups born approximately 90% of wild type size [17]. In contrast, *Igf2r* KO pups had an overgrowth phenotype and were at birth around 140% the mass of wild types [18–20]. The growth restricted phenotypes of either *Igf1* KO or *Igf1r* KO pups were seen only in homozygous mutant offspring, in keeping with equivalent expression of these genes from both their respective maternal and paternal alleles. Importantly, the growth restricted phenotype of heterozygous pups inheriting a mutant paternal allele (*Igf2^+/p^*) was indistinguishable from that of homozygous null (*Igf2^-/-^*) animals, while heterozygous pups inheriting the same allele from their mother (*Igf2^m/+^*) were indistinguishable from wild types (*Igf2^+/+^*) [14]. Similarly, both *Igf2r^m/+^* and *Igf2r^-/-^*pups exhibited a characteristic overgrowth phenotype compared to *Igf2r^+/p^* and *Igf2r^+/+^* animals [19, 20]. Thus, mouse KO experiments established the functional imprinting status of *Igf2* and *Igf2r*, as well as their roles in fetal growth regulation. The importance of IGF signalling for human fetal development is exemplified by characteristic overgrowth in Beckwith-Wiedemann syndrome (BWS), associated with excess *IGF2* expression, and growth restriction in Silver-Russell syndrome (SRS) associated with loss of *IGF2* expression [11, 12, 21].

Epistatic tests involving crosses between different mutant mouse strains demonstrated unequivocally the fetal growth regulatory interactions between the Ins/IGF ligands and receptors (Figure 1A). *Igf1^-/-^*: *Igf1r^-/-^* double knockout (DKO) pups were born approximately 45% of wild type mass and indistinguishable from *Igf1^-/-^* pups, indicating that Igf1 stimulates fetal growth exclusively through Igf1r [15]. In contrast, *Igf2^+/p^*: *Igf1^-/-^* DKO pups were smaller, at around 30% of wild type, indicating an interaction between Igf2 and Igf1r but also an additional receptor [15]. Further epistatic tests ruled out the Insr as the additional receptor [17], despite that Igf2 can bind the Insr, albeit with low affinity (reviewed in [22]). In contrast, *Igf2r^m/+^*pups were large at birth (125-130% relative to wild type), with elevated serum Igf2 levels [18–20]. This overgrowth phenotype was abrogated in *Igf2^+/p^*: *Igf2r^m/+^*DKO mice which, like *Igf2^+/p^* single mutants, were only 60% in size [19, 20, 23]. These data were consistent with the other known role of Igf2r (synonym: cation-independent mannose-6-phosphate receptor) as a receptor that binds mannose-6-phosphate- tagged proteins and internalizes them for lysosomal degradation [24]. The binding site for Igf2 is separate from that for mannose-6-phosphate and is not conserved in the equivalent receptor of non-mammalian vertebrates such as chicken and zebrafish [25, 26], supporting the idea that Igf2 binding by Igf2r represents a step in the evolution of imprinted growth regulation. In summary, it is clear that in mammals *Igf2* and *Igf2r* are oppositely imprinted genes that have opposing influences on fetal growth that fit with the parent-offspring conflict hypothesis [27].

Growth factor receptor-bound protein 10 (Grb10) is a signaling adaptor protein, capable of interacting with numerous different receptor tyrosine kinases (RTKs), typically inhibiting receptor activity and downstream signalling [28–30], in at least some cases through a mechanism involving phosphorylation of Grb10 by raptor kinase, a component of the mTORC1 complex [31–33]. *Grb10* is unusual among imprinted genes in being expressed predominantly from the paternal allele in the developing and adult central nervous system (CNS), and from the maternal allele in tissues outside of the CNS [34–36]. Mice with a germline knockout of the maternal *Grb10* allele (*Grb10^m/+^*) are at birth around 30% larger by weight than wild type littermates [36–38], establishing a role for maternal *Grb10* as a potent inhibitor of fetal growth. While the mass of organs such as lungs and heart increased roughly in line with the whole body, brain size did not increase significantly and was small relative to the body in *Grb10* KO pups. This correlates with the lack of expression from the *Grb10* maternal allele in the CNS, though interestingly no obvious effect on brain size at birth was seen in *Grb10^+/p^* pups and instead paternal *Grb10* expression in CNS has been associated with specific behavioural changes [36, 39–41]. In contrast to brain, *Grb10^m/+^* liver mass was at birth over twice that of wild type littermates [36–38]. This disproportionate enlargement was associated with excessive accumulation of lipid by hepatocytes whereas generally the excess growth involved changes in cell cycle and increased cell number during fetal development [38, 42]. Notably, skeletal muscle mass was increased at birth due to an increase in myofiber number, without changes in myofiber size or in the ratio of fast- and slow-twitch fibres [43], and this increase in muscle or lean mass persists into adulthood [38, 43–45].

Mice overexpressing *Grb10,* due to deletion of imprinting control regions that normally suppress expression of the paternally inherited allele, are born small (around 60% the mass of wild type littermates) and remain small into adulthood, modelling the situation in around 10-20% of growth restricted SRS patients who inherit two maternal copies of the chromosome 7 region containing *GRB10* [12, 21]. This illustrates a conserved role for *GRB10* in fetal growth control that is emphasized by genome-wide association studies (GWAS) in which *GRB10* has been linked with birth weight or body size in several mammalian populations, including human [46], pig [47], sheep [48]and Arctic ringed seal [49].

Mouse studies have shown that Grb10 regulates the Insr *in vivo* to influence glucose regulation through actions on peripheral tissues [33, 44, 50] and the endocrine pancreas [51], and are consistent with human population studies linking *GRB10* with energy homeostasis and endocrine pancreas function (e.g. [52]). Grb10 has also been shown to inhibit Igf1r activity in adult tissues [44, 51] and it is widely assumed that Grb10 influences fetal growth by acting on the Igf1r (Figure 1B). We previously tested this assumption by performing crosses between *Grb10* KO and *Igf2* KO mouse mutants [37]. Resulting *Grb10^m/+^*:*Igf2^+/p^*DKO pups were intermediate in size at birth, compared to *Grb10^m/+^* (large) and *Igf2^+/p^* (small) pups, indicating additive effects of two growth regulators acting largely independently of each other. Since Igf2 influences fetal growth through the Igf1r, together with Igf1 [15–17] (Figure 1A) these experiments formed only an indirect assessment of the potential for Grb10 to act via Igf1r. Given the unexpected nature of this result and the potential for some form of compensation occurring at the level of the receptor, here we tested directly for genetic interactions between *Grb10* and either *Igf1r* or *Insr*. We present two key findings. First, our data support the conclusion that Grb10 regulates fetal growth largely through acting independently of Igf1r or Insr signaling. Second, excessive lipid accumulation in the neonatal *Grb10^m/+^* liver was found to be *Insr*-dependent, meaning that Grb10 modulation of *Insr*-regulated metabolism begins during fetal development. These findings are important for the understanding of fetal growth regulation and its impact on tissue proportions and life-long metabolic health.

## Materials and Methods

### Mice

Generation of the mouse strains *Grb10Δ2-4* (full designation *Grb10^Gt(β-geo)1Ward^*) and *Grb10ins7* (previously referred to as *Grb10KO*; full designation *Grb10^Gt(β-geo)2Ward^*) from gene-trap embryonic stem cell lines has previously been described [36, 37]. Both lines are predicted null alleles and contain a functional *LacZ* reporter gene insertion expressed under the control of endogenous *Grb10* regulatory elements. Detailed characterisation has shown that in *Grb10Δ2-4* the *LacZ* reporter gene has replaced some 36kb of endogenous sequence, including the first 3 protein coding exons (exons 2-4), while the *Grb10ins7* insertion site is associated with a 12bp deletion at the 3’ end of exon 7 [53]. The *Insr KO* [54] and *Igf1r KO* [55] strains, again null alleles, have also been described. To generate experimental animals, first *Grb10Δ2-4^+/p^* and *Grb10ins7^+/p^* males were each crossed with *Igf1r^+/-^* females to generate double heterozygous animals, *Grb10Δ2-4^+/p^*: *Igf1r^+/-^* and *Grb10ins7^+/p^*: *Igf1r^+/-^*. Double heterozygous females were then crossed with *Grb10^+/+^*:*Igf1r^+/-^*males to produce offspring of six genotypes (Table 1a). Mice were genotyped by PCR using primers and conditions previously described for *Grb10* [53] and *Igf1r* [55].

**Table 1.**
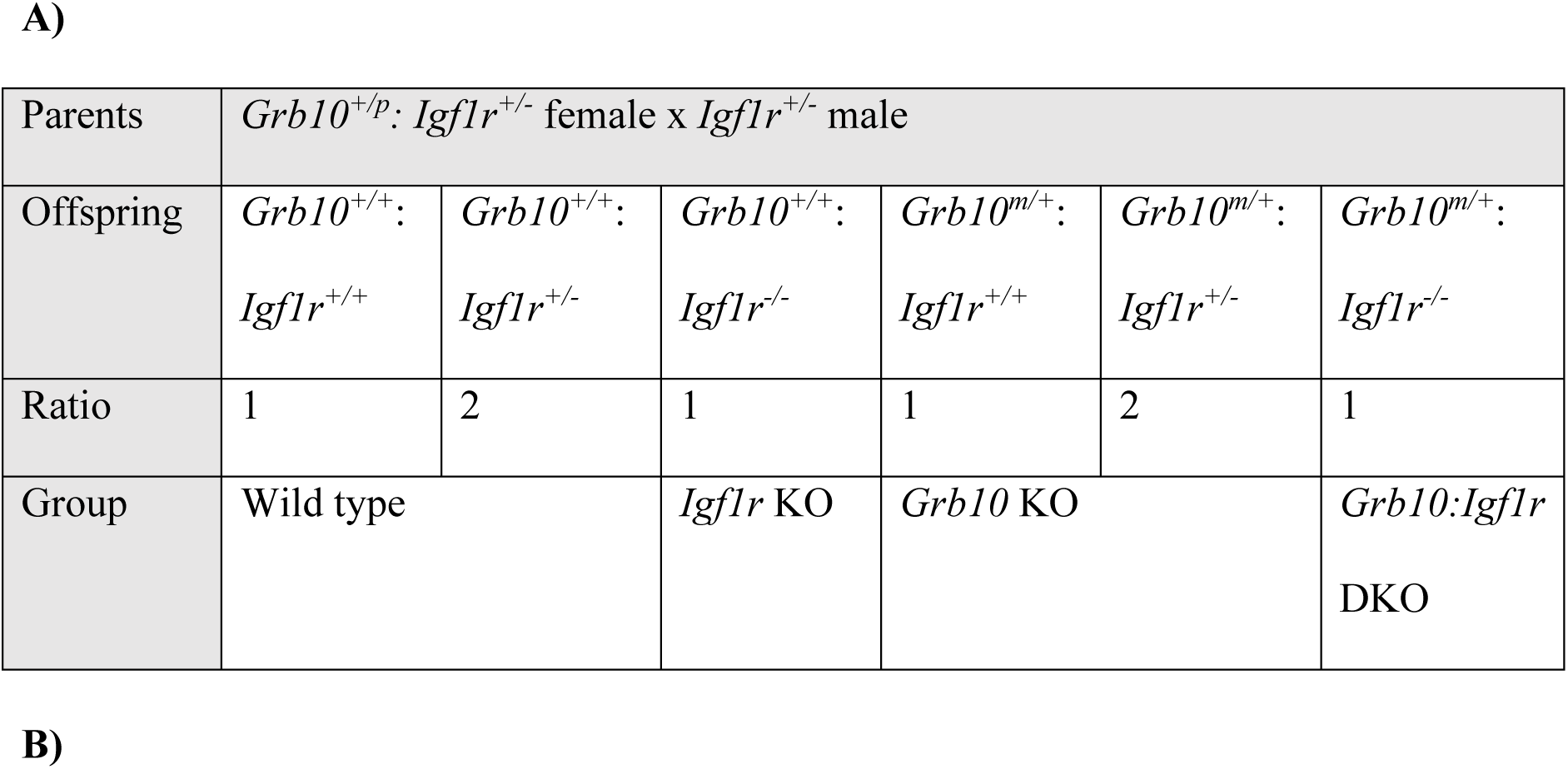
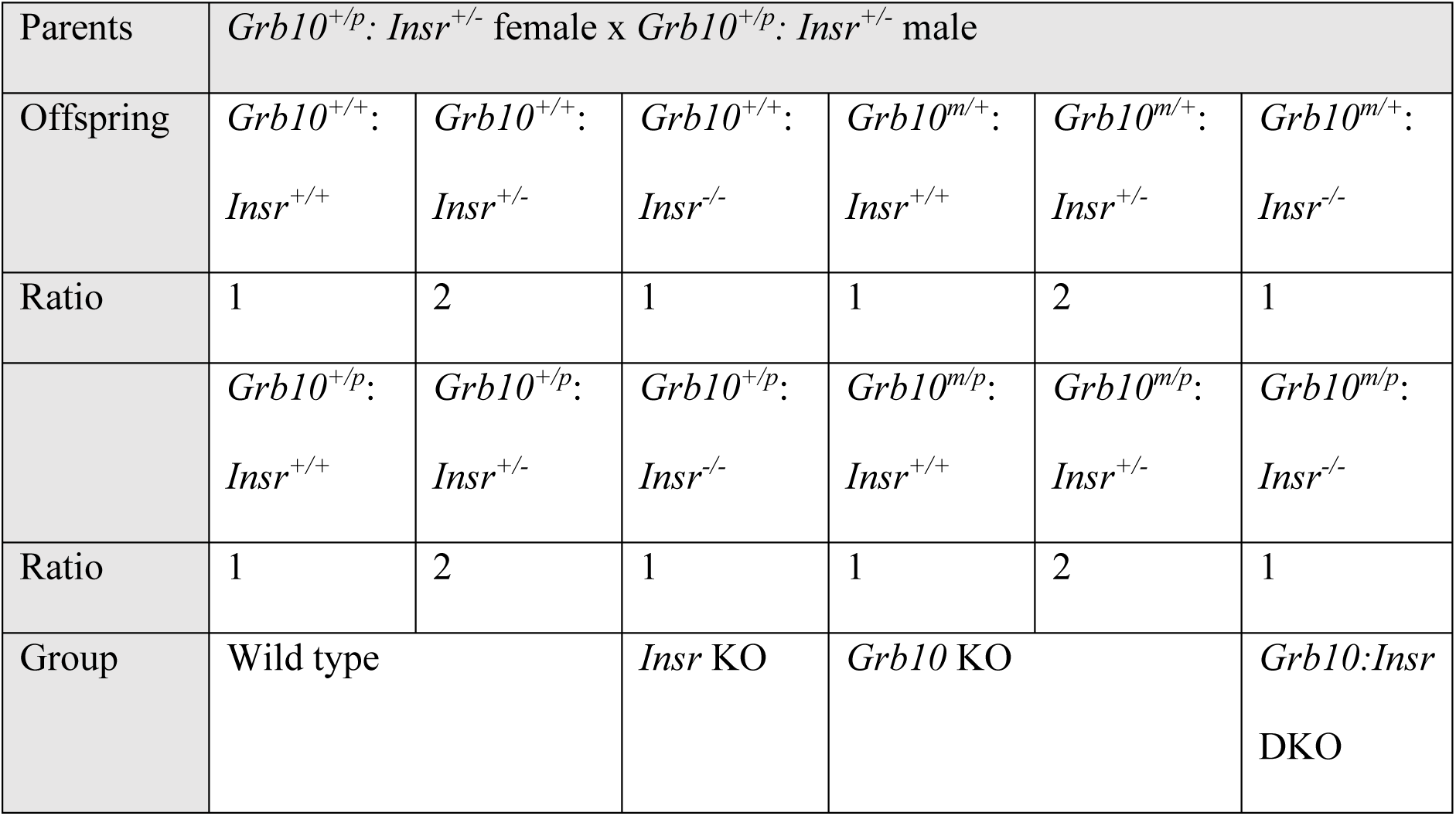
Genetic crosses used in the study, showing parent and offspring genotypes with their expected Mendelian ratios. A) crosses between either *Grb10* KO strains, (*Grb10Δ2-4* and *Grb10ins7*) and the *Igf1r* KO strain. For statistical analysis *Igf1r^+/-^* heterozygous offspring were grouped with their respective *Igf1r^+/+^* wild type counterparts, as indicated. B) Crosses between the *Grb10Δ2-4* KO and *Insr* KO strains. For statistical analysis *Insr^+/-^*heterozygous offspring were grouped with their respective *Insr^+/+^* wild type counterparts, *Grb10^+/p^*with respective *Grb10^+/+^* wild types and *Grb10^m/p^*with respective *Grb10^m/+^* as *Grb10* maternal allele knockouts, as indicated. DKO = double knockout.

*Grb10Δ2-4^+/p^* males were also crossed with *Insr^+/-^*females to generate double heterozygous animals, *Grb10Δ2-4^+/p^*: *Insr^+/-^*. These double heterozygous females were intercrossed to produce offspring of 12 genotypes (Table 1b). In addition to using PCR to genotype offspring for wild type and mutant *Grb10Δ2-4* [53] and *Insr* [17] alleles, carcasses were *LacZ* stained [36] to determine the parental origin of mutant *Grb10* alleles. Embryos and placentae were collected on embryonic day e17.5, where e0.5 was the day on which a copulation plug was observed. Otherwise, experimental animals were collected on the day of birth, designated post-natal day 1 (PN1). Wild type littermates are considered the control group and single animals the biological replicate, noting that multiple litters were generated in each cross, with the aim of having enough of the least common genotypes for robust statistical analysis. All animals were maintained on a mixed inbred (C57BL/6J:CBA/Ca) strain background and housed under conditions of 13 hours light:11 hours darkness, including 30-minute periods of dim lighting to provide false dawn and dusk, a temperature of 21±2°C and relative humidity of 55±10%. Standard chow (CRM formula; Special Diets Services, Witham, Essex, UK) and water was freely available.

### Tissue collection, histology and blood glucose measurements

Whole bodies and organs were collected, any surface fluid removed from embryos or dissected organs by gently touching them onto absorbent paper, and weights obtained using a fine balance accurate to 4 decimal places (Sartorius BP61S). Paired organs (lungs and kidneys) were weighed together. Organs for histology were fixed by immersion in 4% (w/v) paraformaldehyde (PFA) in PBS at 4°C for 16-24 hours, then processed by machine (Leica TP1020) for wax embedding. Sections were cut at approximately 8-10 μm using a microtome (Leica Histocore Biocut), prior to staining with haematoxylin and eosin (H & E) as previously described [56]. Images were collected using a digital colour camera (Olympus SC50) and software (Olympus cellSens Entry), attached to a compound microscope (Nikon Eclipse E800), then scored with the operator blind to genotype. Glucose measurements were obtained using a One-Touch ULTRA (Lifescan, CA) glucometer immediately following collection of whole blood by decapitation of PN1 pups.

### Statistical analysis

Chi-square tests were applied to determine whether the genotypes of experimental groups were present in the expected Mendelian ratios. Otherwise, numerical data were subject to one-way analysis of variance (ANOVA), using a Kruskall-Wallis test with post-hoc Dunn’s test to determine p-values between groups. This test allowed us to detect significant differences associated with either of the single knockout groups in each set of progeny as well as any significant interaction between them. This relatively conservative non-parametric test was chosen because in some experiments one or more genotype group was represented by a small samples size (n=<5). In order to test for an interaction between mutant genotypes we also applied a two-way ANOVA test where indicated. All statistical tests were applied using GraphPad Prism (v10 GraphPad, La Jolla, CA, USA) software.

Graphs show arithmetic means ±standard error of the mean (SEM). Differences with p-values of <0.05 were considered statistically significant.

## Results

### Analysis of genetic interaction between *Grb10* and *Igf1r* for the regulation of fetal growth

To directly assess the possibility that *Grb10* interacts with the *Igf1r* to influence growth we performed genetic crosses between both *Grb10Δ2-4* and *Grb10ins7* (collectively referred to as *Grb10* KO strains) and *Igf1r* KO mice. *Grb10Δ2-4* offspring were analysed at PN1 and e17.5 whereas *Grb10ins7* offspring were analysed at PN1 only. To increase statistical power, both sexes were pooled together and considered in a single analysis, with mean weights ± standard error of the mean stated in the text and shown graphically for offspring genotype groups. PN1 data were consistent between offspring of the two *Grb10* KO strains (as summarised in Table 2) and consequently all subsequent experiments were carried out with only the *Grb10Δ2-4* strain.

**Table 2.** Summary of PN1body and organ weight data for progeny of crosses between *Grb10* KO strains and *Igf1r* KO mice. Mean weights are shown for each genotype together with changes relative to wild type (%WT) for each mutant genotype. A) *Grb10ins7* KO data. B) *Grb10Δ2-4* KO data.

|  | WT | *Igf1r* KO |  | *Grb10* KO |  | DKO |  |
| --- | --- | --- | --- | --- | --- | --- | --- |
|  | Actual | Actual | % WT | Actual | % WT | Actual | % WT |
| Body | 1.4010 | 0.6395 | -54 | 1.7670 | +26 | 1.1650 | -17 |
| Brain | 0.0824 | 0.0491 | -40 | 0.0860 | +4 | 0.0535 | -35 |
| Liver | 0.0550 | 0.0469 | -15 | 0.1231 | +124 | 0.1115 | +103 |
| Lung | 0.0387 | 0.0079 | -80 | 0.0480 | +24 | 0.0287 | -26 |
| Heart | 0.0092 | 0.0078 | -15 | 0.0127 | +39 | 0.0094 | +2 |
| Kidney | 0.0159 | 0.0111 | -31 | 0.0166 | +5% | 0.0128 | -20 |

**Table 2.** Summary of PN1 body and organ weight data for progeny of crosses between *Grb10* KO strains and *Igf1r* KO mice. Mean weights are shown for each genotype together with changes relative to wild type (%WT) for each mutant genotype. A) *Grb10ins7* KO data. B) *Grb10 Δ2-4* KO data.
|  | WT | *Igf1r* KO |  | *Grb10Δ2-4* KO |  | DKO |  |
| --- | --- | --- | --- | --- | --- | --- | --- |
|  | Actual | Actual | % WT | Actual | % WT | Actual | % WT |
| Body | 1.434 | 0.5984 | -58 | 1.9340 | +35 | 1.3620 | -5 |
| Brain | 0.105 | 0.0499 | -48 | 0.0958 | -9 | 0.0603 | -43 |
| Liver | 0.0579 | 0.0444 | -23 | 0.1297 | +124 | 0.1322 | +128 |
| Lung | 0.0404 | 0.0087 | -78 | 0.0460 | +14 | 0.0353 | -13 |
| Heart | 0.0088 | 0.0071 | -20 | 0.0158 | +79 | 0.0120 | +36 |
| Kidney | 0.0239 | 0.0091 | -62 | 0.0183 | -23% | 0.0143 | -40 |

### *Grb10ins7* KO x *Igf1r* KO offspring PN1 body mass

Progeny of crosses between *Grb10ins7^+/p^*:*Igf1r^+/-^*females and *Grb10ins7^+/+^*:*Igf1r^+/-^* males were collected at PN1 for body and organ weight analysis (Figure 2). Progeny with six genotypes were reduced to four groups by pooling *Grb10ins7^+/+^:Igf1r^+/-^*with *Grb10ins7^+/+^:Igf1r^+/+^* (wild type group) and *Grb10ins7^m/+^:Igf1r^+/-^* and *Grb10in7^m/+^:Igf1r^+/+^*(*Grb10ins7* KO group), for comparison with the *Igf1r* KO and *Grb10ins7*:*Igf1r* DKO groups (Table 1a). This was done following initial analysis of the data which confirmed that *Igf1r^+/-^*animals had a normal fetal growth phenotype, as previously shown [15]. Pooling allowed us to strengthen statistical analyses, while simplifying data analysis and presentation, without materially affecting the outcome. If Grb10 regulates growth through an interaction with the Igf1r, *Grb10*:*Igf1r* DKO animals would be expected to be phenotypically indistinguishable from *Igf1r* KO animals (Figure 1B). Body mass data (Figure 2A, Table 2A) immediately indicated that we should reject this hypothesis. *Grb10ins7* KO pups (mean weight 1.7670±0.0360g) were approximately 26% larger (p<0.0001) and *Igf1r* KOs (0.6395±0.0267g) 54% smaller (p<0.01) than wild type controls (1.401±0.0297g), respectively, whereas *Grb10ins7:Igf1r* DKO mutants were intermediate in size (1.1650±0.0554g). Thus, *Grb10ins7*:*Igf1r* DKO pups displayed an additive effect of both parental genotypes, being significantly different from both *Igf1r* KO (p<0.01) and *Grb10ins7* KO (p<0.0001) single mutants, but not from wild type neonates (Figure 2A). This was supported by a two-way ANOVA test which detected no interaction between the two genotypes (p=0.1017).

**Figure 2.**
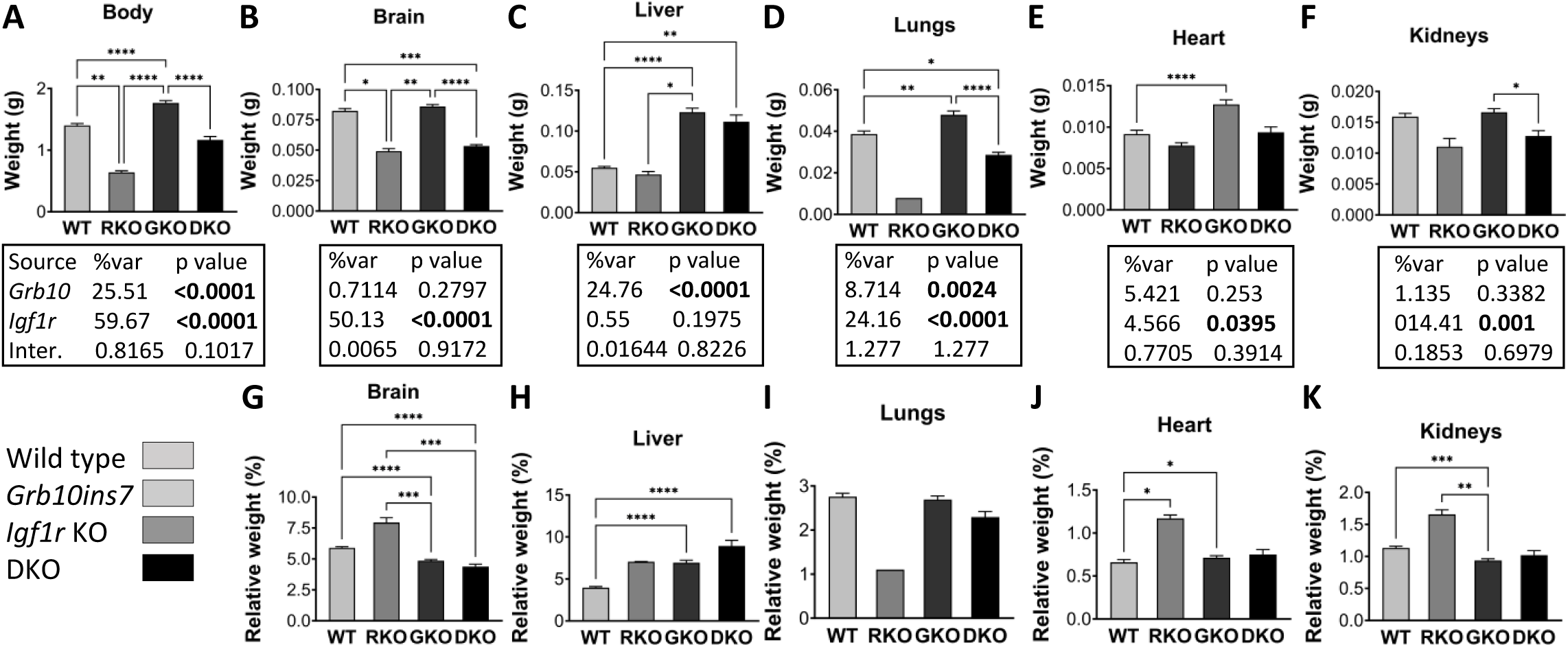
Weights of whole body and selected dissected organs at PN1 from progeny of crosses between *Grb10ins7* KO and *Igf1r* KO mice. Data were pooled into four groups for analysis as described in the Methods, wild type (WT), *Igf1r* KO (RKO), *Grb10* KO (GKO) and *Grb10*:*Igf1r* double knockouts (DKO). Body weights are shown for the four offspring genotype groups (A). Actual weights of brain (B), liver (C), lungs (D), heart (E) and kidneys (F) are shown alongside relative weights of the same organs, expressed as a percentage of body mass (G-K). Values represent means and SEM, tested by one-way ANOVA using Kruskal-Wallis and Dunn’s post hoc statistical tests. Summaries of Two- way ANOVA outcomes beneath each graph show the percentage of total variation (%var) and a p value for each source, namely the two single KO genotypes and any interaction (Inter.) between the two (values significant at p<0.05 in bold). Sample sizes were, for body, WT n=38, *Igf1r* KO n=7, *Grb10* KO n=26, *Grb10*:*Igf1r* DKO n=12; brain, WT n=38, *Igf1r* KO n=3, *Grb10* KO n=25, *Grb10*:*Igf1r* DKO n=8; liver, WT n=38, *Igf1r* KO n=2, *Grb10* KO n=25, *Grb10*:*Igf1r* DKO n=7; lungs, WT n=38, *Igf1r* KO n=7, *Grb10* KO n=12, *Grb10*:*Igf1r* DKO n=7; heart, WT n=37, *Igf1r* KO n=2, *Grb10* KO n=8, *Grb10*:*Igf1r* DKO n=7; kidneys, WT n=38, *Igf1r* KO n=2, *Grb10* KO n=25, *Grb10*:*Igf1r* DKO n=7. Asterisks indicate *p*-values, *p <0.05, **p <0.01, ***p <0.001, ****p<0.0001.

### *Grb10ins7* KO x *Igf1r* KO offspring PN1 organ mass

To assess body proportions selected individual organs (brain, liver, lungs, heart, kidneys) were dissected at PN1 and their weights were analysed directly (Figure 2B-F) and as a percentage of total body weight (Figure 2G-K). The pattern of organ weight difference across the genotypes was again consistent with the DKO pups having an additive phenotype, comprising the sum of the two single KO phenotypes (summarised in Table 2A). First, the brain from *Grb10ins7* KO (mean mass 0.0860±0.0017g) pups was spared from the general overgrowth phenotype indicated by body mass and was only 4% larger than wild type brain (0.0824±0.0019g) (Figure 2B). Meanwhile, brains from *Igf1r* KO (0.0491±0.0021g) and *Grb10ins7*:*Igf1r* DKO (0.0535±0.0012g) pups were strikingly similar, being smaller than wild type brain by 40% (p<0.05 ), and 35% (p<0.001), respectively. Thus, while *Igf1r* KO brains were roughly proportionate with body size, both *Grb10ins7* KO (p<0.0001) and *Grb10ins7*:*Igf1r* DKO (p<0.001) brains were disproportionately small within larger bodies (Figure 2G). In other words, the *Grb10ins7*:*Igf1r* DKO phenotype was dominated by brain size being severely reduced, as in *Igf1r* KO pups, which can therefore be attributed to loss of *Igf1r* expression.

In direct contrast, the livers of *Grb10ins7* KO (0.1231±0.0051g) and *Grb10ins7*:*Igf1r* DKO (0.1115±0.0083g) pups were each at least double, by 124% (p<0.0001) and 103% (p0.01), respectively, the size of wild type (0.0550±0.0016g), while the *Igf1r* KO (0.0469±0.0036g) liver was some 15% smaller. (Figure 2C). Consequently, while the liver was disproportionately enlarged within the heavier *Grb10ins7* KO body (p<0.0001), liver disproportion was exaggerated in *Grb10ins7*:*Igf1r* DKO (p<0.0001) pups, due to DKOs having a body size similar to wild type (Figure 2H). Due to their greatly reduced body mass relative to wild types, although *Igf1r* KO livers were smaller in actual mass than in wild type controls, *Igf1r* KO pups also had disproportionately large livers. Thus, the *Grb10ins7*:*Igf1r* DKO liver weight phenotype was clearly dominated by the massive size increase also seen in *Grb10* KO single mutants and therefore associated with loss of the maternal *Grb10ins7* allele.

The remaining organs followed a pattern of size difference like that seen in the body mass data, in that *Grb10ins7*:*Igf1r* DKO mass was intermediate between that of the two single KO values. Compared to wild type (0.0387±0.0014g) lungs from a single *Igf1r* KO sample (0.0079g) were 80% lighter (not statistically significant due to very small samples size) and *Grb10ins7* KO (0.0480±0.0019g) 24% heavier (p<0.01), whereas *Grb10ins7*:*Igf1r* DKO (0.0287±0.0013g) lungs were 26% smaller (p<0.05) and intermediate in size (Figure 2D). Relative to total body mass, *Igf1r* KO lungs appeared disproportionately small while *Grb10ins7* KO and *Grb10ins7*:*Igf1r* DKO lungs were roughly proportionate with their respective body sizes (Figure 2I). Similarly, in comparison with wild type (0.0092±0.0005g), hearts from *Igf1r* KO (0.0078±0.0004g) pups were some 15% smaller and *Grb10ins7* KO hearts (0.0127±0.0006g) 39% larger (p<0.0001), with *Grb10ins7*:*Igf1r* DKO hearts (0.0094±0.0006g) intermediate in size, being only 2% larger than wild type (Figure 2E). While these weight differences were not all statistically significant, in relative terms, the heart from *Grb10ins7* KO (p<0.05) and *Igf1* KO (p<0.05) single mutants were disproportionately large, whereas the *Grb10ins7*:*Igf1r* DKO heart was not (Figure 2J).

In the case of kidneys, those from *Grb10ins7* KO (0.0166±0.0006g) were only slightly enlarged, by 5%, compared with wild type (0.0159±0.0005g), while both *Igf1r* KO (0.0111±0.00134g) and DKO (0.01287±0.0009g), were smaller by 31% and 20%, respectively (Figure 2F). The only significant difference in kidney weights was between *Grb10ins7* KO and *Grb10ins7*:*Igf1r* DKO (p<0.05). Relative to wild type body mass, this meant that *Grb10ins7* KO pups alone had disproportionately small kidneys (p<0.001) (Figure 2K). For each individual organ two-way ANOVA tests indicated there was no interaction between the genotypes, just as for the whole body (Figure 2A-F).

### *Grb10Δ2-4* KO x *Igf1r* KO offspring PN1 body mass

To corroborate data from the *Grb10ins7* strain, similar PN1 data were collected using the *Grb10Δ2-4* strain. Progeny of crosses between *Grb10Δ2-4^+/p^*:*Igf1r^+/-^* females and *Grb10Δ2-4^+/+^*:*Igf1r^+/-^* males were again collected at PN1 and whole body weights recorded along with weights of selected organs (Figure 3). As before, data for the six offspring genotypes were pooled to generate four groups for analysis, combining *Grb10Δ2- 4^+/+^:Igf1r^+/-^* with *Grb10Δ2-4^+/+^:Igf1r^+/+^* (wild type group) and *Grb10Δ2-4^m/+^:Igf1r^+/-^*with *Grb10Δ2-4^m/+^:Igf1r^+/+^* (*Grb10Δ2-4* KO group) progeny (Table 2B), which was again supported by our initial data analysis. As for the previous cross, while *Grb10Δ2-4* KO pups (mean weight 1.9340 ±0.0283g) were around 35% larger (p<0.0001) and *Igf1r* KOs (0.5984±0.0270g) 58% smaller (p<0.0001), respectively, than wild type controls (1.4340±0.0244g), *Grb10Δ2-4:Igf1r* DKO mutants (1.3620±0.0422g) were intermediate in size, just 5% smaller than wild type (Figure 3A; Supp Figure 2A). *Grb10Δ2-4:Igf1r* DKO pups were significantly smaller than *Grb10Δ2-4* KO pups (p<0.0001) but not smaller than wild type neonates (Figure 3A, B), while *Grb10Δ2-4* KO pups were significantly larger than both wild type (p<0.0001) and *Igf1r* KO (p<0.0001) pups. It should be noted that in this dataset although *Igf1r* KO pups were very small, as expected [15], the difference between them and wild type pups did not reach significance, as it did in the previous cross involving the *Grb10ins7* line, likely because of the small number (n = 6) of *Igf1r* KO pups obtained in this case. In line with this, the two-way ANOVA test indicated a possible interaction between the genotypes, but at a relatively high significance level (p=0.0135). Despite this, it was clear that *Grb10Δ2-4:Igf1r* DKO pups were not small, to the extent consistently shown for *Igf1r* KO pups, and instead their intermediate size must result from an additive effect of the two mutant parental genotypes.

**Figure 3.**
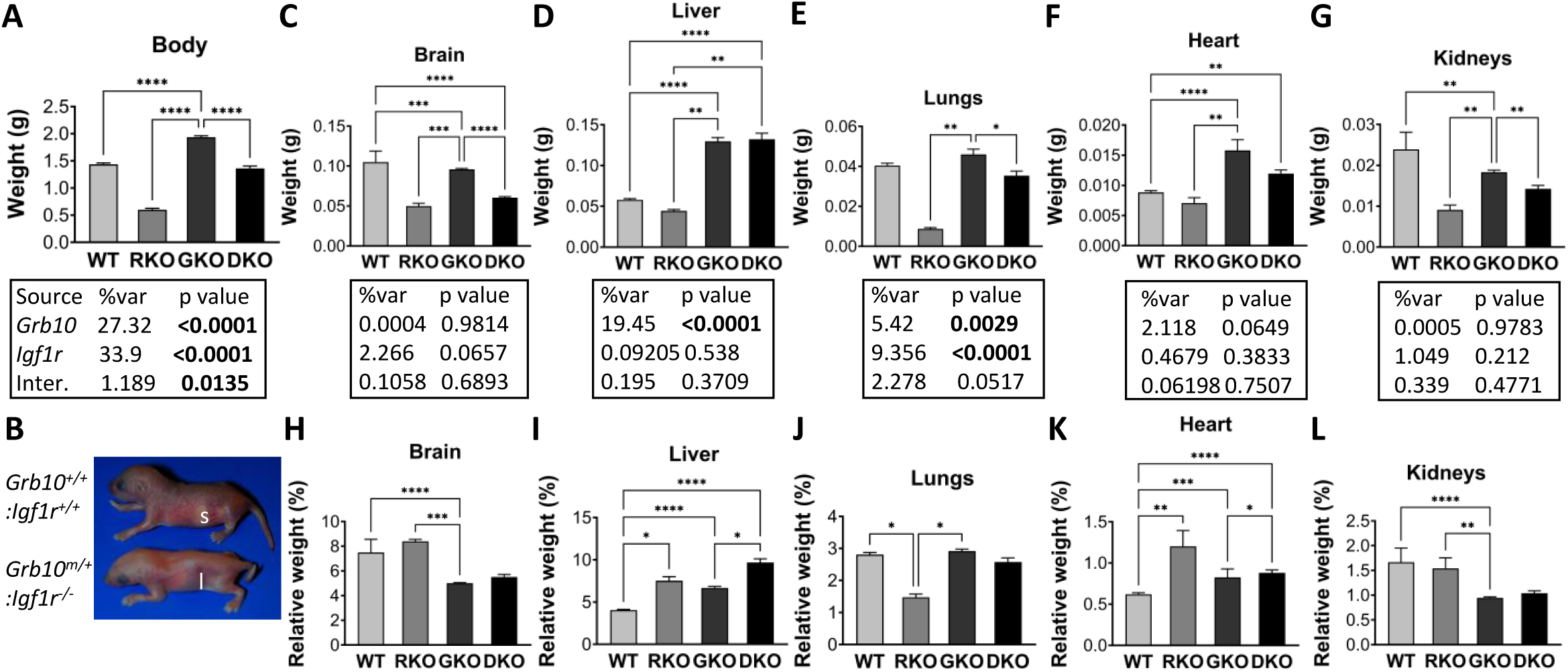
Weights of whole body and selected dissected organs at PN1 from progeny of crosses between *Grb10Δ2-4* KO and *Igf1r* KO mice. Data were pooled into four groups for analysis as described in the Methods, wild type (WT), *Igf1r* KO (RKO), *Grb10* KO (GKO) and *Grb10*:*Igf1r* double knockouts (DKO). Body weights are shown for the four offspring genotype groups (A). Gross physical appearance of typical WT (*Grb10^+/+^*:*Igf1r^+/+^*) and *Grb10*:*Igf1r* DKO (*Grb10^-/-^*:*Igf1r^-/-^*) pups, noting in the DKO the small head relative to body size and enlarged liver (l) obscuring the milk filled stomach (s), clearly visible through the skin of the wild type (B). Actual weights of brain (C), liver (D), lungs (E), heart (F) and kidneys (G) are shown alongside relative weights of the same organs, expressed as a percentage of body mass (H-L). Values represent means and SEM, tested by one-way ANOVA using Kruskal-Wallis and Dunn’s post hoc statistical tests. Summaries of Two-way ANOVA outcomes beneath each graph show the percentage of total variation (%var) and a p value (values significant at p<0.05 in bold) for each source, namely the two single KO genotypes and any interaction (Inter.) between the two. Sample sizes were, for body, WT n=66, *Igf1r* KO n=6, *Grb10* KO n=66, *Grb10*:*Igf1r* DKO n=16; brain, WT n=66, *Igf1r* KO n=3, *Grb10* KO n=65, *Grb10*:*Igf1r* DKO n=16; liver, WT n=66, *Igf1r* KO n=3, *Grb10* KO n=65, *Grb10*:*Igf1r* DKO n=16; lungs, WT n=66, *Igf1r* KO n=3, *Grb10* KO n=66, *Grb10*:*Igf1r* DKO n=16; heart, WT n=66, *Igf1r* KO n=3, *Grb10* KO n=65, *Grb10*:*Igf1r* DKO n=16; kidneys, WT n=66, *Igf1r* KO n=3, *Grb10* KO n=65, *Grb10*:*Igf1r* DKO n=16. Asterisks indicate *p*-values, *p <0.05, **p <0.01, ***p <0.001, ****p<0.0001.

### *Grb10Δ2-4* KO x *Igf1r* KO offspring PN1 organ mass

As before, body proportions were assessed by dissecting and weighing selected organs at PN1. Organ weights were analysed directly (Figure 3C-G) and as a percentage of total body weight (Figure 3H-L). The genotype-dependent differences in organ weights were again consistent with the *Grb10Δ2-4:Igf1r* DKO pups having an additive phenotype in comparison with the two single KO genotypes (summarised in Table 2B). First, the brain from *Grb10Δ2-4* KO (0.0958±0.0011g) pups was spared from the general overgrowth phenotype indicated by body mass and was in fact some 9% smaller (p<0.001) than wild type brain (0.1050±0.01376g) (Figure 3C), Meanwhile, brains from *Igf1r* KO (0.0499±0.0034g) and *Grb10Δ2-4:Igf1r* DKO (0.0603±0.0140g) pups were strikingly similar, being smaller than wild type brain by 52%, and 43% (p<0.001), respectively. Thus, while *Igf1r* KO brains were proportionate to their small bodies, both *Grb10Δ2-4* KO (p<0.0001) and *Grb10Δ2-4:Igf1r* DKO brains were small (not significantly so, likely due to the small number of DKO weighed; n=3) within larger bodies (Figure 3H). In other words, the *Grb10Δ2-4:Igf1r* DKO phenotype was dominated by brain size being severely reduced, as in *Igf1r* KO pups, and is therefore associated with loss of *Igf1r* expression.

In contrast, the livers of DKO (0.1322±0.0075g) and *Grb10Δ2-4* KO (0.1297±0.0045g) pups were again each more than double (128% and 124% larger, respectively) the size of wild type (0.0579±0.0016g) liver (p<0.0001), while the *Igf1r* KO (0.0444±0.0018g) liver was some 23% smaller (Figure 3D). Consequently, while the liver was disproportionately enlarged within the heavier *Grb10Δ2-4* KO body (p<0.0001), in the *Grb10Δ2-4:Igf1r* DKO liver disproportion was exaggerated (p<0.0001), due to DKOs having a body size similar to wild type (Figure 3I). Due to their greatly reduced body mass relative to wild types, although *Igf1r* KO livers were smaller in actual mass than in wild type controls, *Igf1r* KO pups also had disproportionately large livers. Similar to our findings using the *Grb10ins7* KO strain, the *Grb10Δ2-4:Igf1r* DKO phenotype was clearly dominated by the massive size increase associated with loss of the maternal *Grb10Δ2-4* allele.

The remaining organs followed a pattern of size differences like that seen in the body mass data, in that DKO mass was intermediate between that of the two single KO values. Lungs from *Grb10Δ2-4:Igf1r* DKO (0.0353±0.0023g) pups were approximately the same size as wild type (0.0404±0.0012g) but differed to those of both single mutants, with *Igf1r* KO (0.0087±0.0065g) approximately 78% lighter than wild type and *Grb10Δ2-4* KO (0.0460±0.0026g) 14% heavier (Figure 3E). *Grb10Δ2-4* KO lung weight was significantly different to both *Igf1r* KO (p<0.01) and *Grb10Δ2-4:Igf1r* DKO (p<0.05). Relative to total body mass, *Igf1r* KO lungs were disproportionately small (p<0.05) while *Grb10Δ2-4* KO and *Grb10Δ2-4:Igf1r* DKO lungs were roughly proportionate with their respective body sizes (Figure 3J). Similarly, in comparison with wild type (0.0088±0.0003g), hearts from *Igf1r* KO (0.0071±0.0009g) pups were some 20% smaller and *Grb10Δ2-4* KO hearts (0.0158±0.0018g) 79% larger (p<0.0001) (Figure 3F). *Grb10Δ2-4:Igf1r* DKO hearts (0.0120±0.0006g) were intermediate in size, being significantly larger than both wild type (p<0.0001) and *Igf1r* KO (p<0.01) single mutants. In relative terms, the hearts from pups of all three mutant genotypes were disproportionately large compared to wild type controls (Figure 3K).

Compared to wild type kidneys (0.0239±0.0042g), *Igf1r* KO (0.0091±0.0012g) and *Grb10Δ2-4:Igf1r* DKO (0.0143±0.0008g) (Figure 3G) kidneys were reduced in size, by 62% and 40%. It should be noted that *Grb10Δ2-4* KO kidneys (0.0183±0.0005g) were also smaller than wild type, by 23% (p<0.01) in absolute terms, therefore exhibiting sparing from the general overgrowth phenotype. Notably, *Grb10Δ2-4* KO kidney weights were still significantly different to those of *Igf1r* KO (p<0.01) and *Grb10Δ2-4:Igf1r* DKO (p<0.01). Relative to body mass (Figure 3L), this meant kidneys were proportionate in the small body of *Igf1r* KO pups, but disproportionately small in the larger body of *Grb10Δ2-4* KO (p<0.0001) pups. As in the previous cross, two-way ANOVA tests for individual organs indicated there was no interaction between the genotypes in each case (Figure 3C-G).

The organ disproportion evident in *Grb10Δ2-4:Igf1r* DKO PN1 pups was reflected by their appearance (Figure 3B). Despite being similar in size to wild types, *Grb10Δ2-4:Igf1r* DKO pups had small, flattened heads and livers that were distended such that they largely obscured the milk-filled stomach.

### *Grb10Δ2-4* KO x *Igf1r* KO offspring e17.5 embryo and placenta

To investigate the potential for interaction between *Igf1r* and *Grb10* to regulate growth by acting within the placenta we analysed weights of the whole embryo and placenta at e17.5 (Figure 4). We chose a time-point late in gestation when any size differences between conceptuses of different genotypes would be relatively large. The pattern of size differences observed was very similar to that seen for pups at PN1. *Grb10Δ2-4* KO embryos (1.085±0.0450g) were 35% larger than wild type (0.8031±0.0371g), whereas the single *Igf1r* KO (0.4029g) embryo collected was 50% smaller and *Grb10Δ2-4:Igf1r* DKO embryos (0.6330±0.0286g) intermediate in size, at 21% lighter than wild types (Figure 4A). Unsurprisingly, the one *Igf1r* KO embryo showed no statistical differences in size compared to any of the other genotypes, however, *Grb10Δ2-4* KO embryos were significantly larger than wild type (p<0.05) and *Grb10Δ2-4:Igf1r* DKO (p<0.0001) embryos.

**Figure 4.**
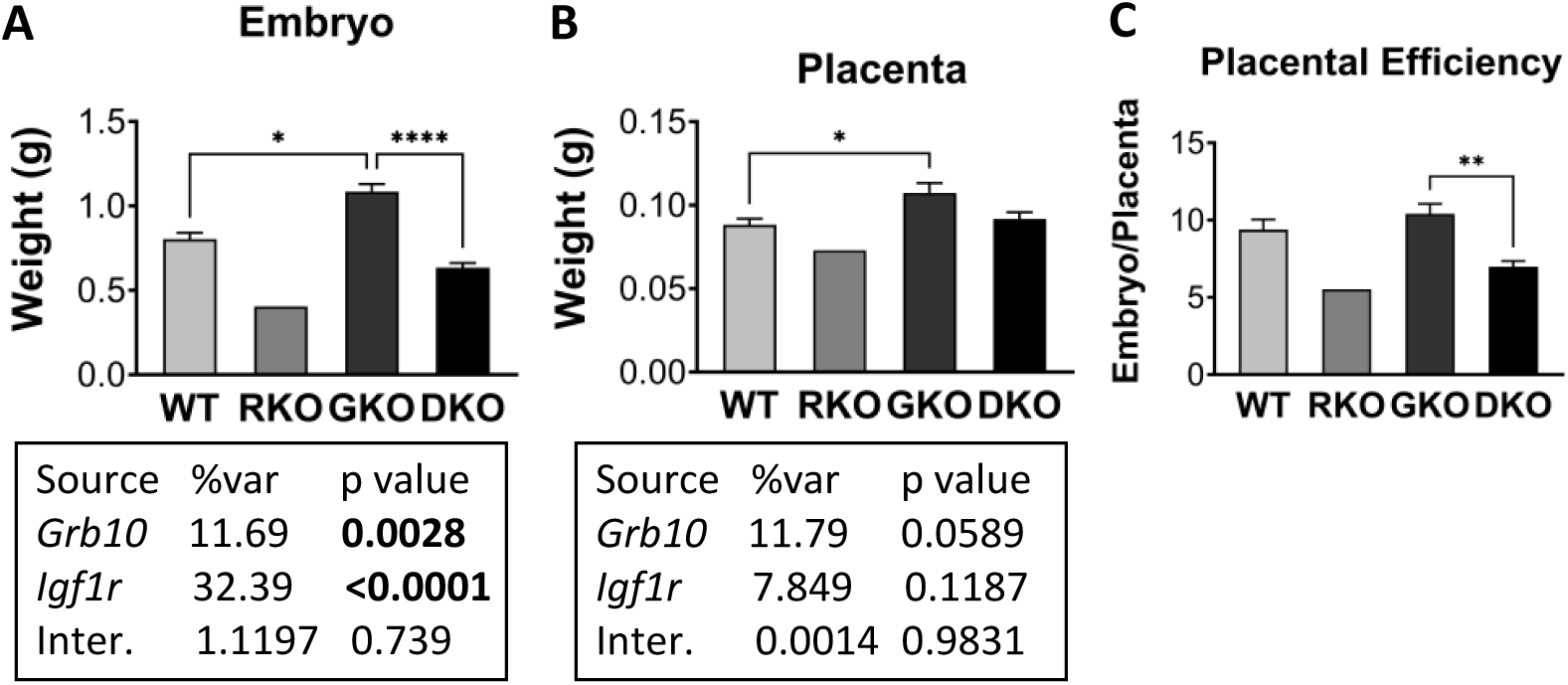
Weight analysis of e17.5 conceptuses from crosses between *Grb10Δ2-4* KO and *Igf1r* KO mice. Data were pooled into four groups for analysis as described in the Methods, wild type (WT), *Igf1r* KO (RKO), *Grb10* KO (GKO) and *Grb10*:*Igf1r* double knockouts (DKO). Weights are shown for the four offspring genotype groups for embryo (A) and placenta (B) and these have been used to calculate the embryo:placenta weight ratio as a measure of placental efficiency (C). Values represent means and SEM, tested by one-way ANOVA using Kruskal-Wallis and Dunn’s post hoc statistical tests. Summaries of Two-way ANOVA outcomes beneath each graph show the percentage of total variation (%var) and a p value (values significant at p<0.05 in bold) for each source, namely the two single KO genotypes and any interaction (Inter.) between the two. Sample sizes were, for WT n=17, *Igf1r* KO n=1, *Grb10* KO n=11, *Grb10*:*Igf1r* DKO n=9. Asterisks indicate *p*- values, *p <0.05, **p <0.01, ****p<0.0001.

Placental weights followed a similar pattern (Figure 4B), with *Grb10Δ2-4* KO (0.1073±0.0060g) 22% larger than wild type (0.0882±0.0036g), the single *Igf1r* KO placenta (0.0729g) 17% smaller and *Grb10Δ2-4:Igf1r* DKO placentas (0.0916±0.0040g) in between at only 4% larger. The only statistically significant difference was between wild type and *Grb10Δ2-4* KO samples (p<0.05). Next, the ratio of embryo to placental mass was calculated for each genotype as an estimate of placental efficiency (Figure 4C).

Although not statistically significant, the trend was for *Grb10Δ2-4* KO placental efficiency (10.41) to be slightly higher than wild type (9.39), while both *Igf1r* KO (5.53) and *Grb10Δ2-4:Igf1r* DKO (6.98) were lower than wild type, with the only significant difference between *Grb10Δ2-4* KO and *Grb10Δ2-4:Igf1r* DKO (p<0.01). A two-way ANOVA test found no evidence of an interaction between the genotypes for either embryo or placenta size (Figure 4A,B).

### Survival of *Grb10* KO x *Igf1r* KO progeny at PN1 and e17.5

During collection of offspring the small, presumptive *Igf1r* KO pups seemed scarce and chi-square tests of observed versus expected numbers generally supported this notion (Supp. Table 1). Testing of PN1 data from the *Grb10Δ2-4* KO x *Igf1r* KO cross, which had the largest sample size (n=154), indicated that paucity of *Igf1r* KO pups was statistically significant (p=0.0139; Supp. Table 1A), with around one third of the expected numbers surviving. The same was true for offspring collected from the same cross at e17.5 (p=0.015), though in this case the sample size was lower (n=38) and only one *Igf1r KO* embryo was obtained, with the expected number being closer to 5 (Supp. Table 2B). In the case of the *Grb10ins7* x *Igf1r* KO PN1 dataset (n=83), the lack of *Igf1r* KO pups was less evident (7 collected from some 10 expected) and the chi-square test indicated no significant deviation from expected mendelian ratios (p=0.4711; Supp. Table 1C). In both crosses it was clear that *Igf1r* KO pups found alive on the day of birth were failing to thrive, as previously reported[16]. Strikingly, this did not appear to be true for *Grb10*:*Igf1r* DKO PN1 pups in either cross which typically had milk-filled stomachs and appeared to be doing well on PN1.

### Genetic analysis of interaction between *Grb10* and *Insr* for the regulation of fetal growth

#### *Grb10Δ2-4* KO x *Insr* KO offspring PN1 body mass

To address the question of whether *Grb10* regulates growth *in vivo* through an interaction with the *Insr*, we next performed intercrosses between *Grb10Δ2-4*^+/p^:*Insr^+/-^* double heterozygous mice, giving rise to twelve offspring genotypes, which were reduced to four groups for analysis (Table 1B). In addition to combining animals with *Insr^+/-^* and *Insr^+/+^* genotypes (*Insr* wild type groups), we also pooled *Grb10Δ2-4*^+/+^ with *Grb10Δ2-4*^+/p^ genotypes (*Grb10* wild type) and *Grb10Δ2-4*^m/+^ with *Grb10Δ2-4*^m/p^ (*Grb10* KO). This is because the *Grb10* paternal allele is silent in the majority of tissues and its knockout is well established to have no effect on fetal growth [36–38, 57]. Similarly, only *Insr^-/-^*animals have been shown to have a mutant phenotype affecting either growth or glucose regulation [17, 54]. Initial analysis of our data prior to pooling was in line with these earlier studies. As for the Igf1r, should Grb10 regulate growth through an interaction with the Insr, *Grb10Δ2-4*:*Insr* DKO animals would be phenotypically indistinguishable from *Insr* KO single mutants (Figure 1B).

Progeny were first collected at PN1 for body and organ weight analysis (Figure 5). As for the crosses involving KOs, body mass data (Figure 5A, Table 3) indicated that we should reject this hypothesis. *Insr* KO pups (1.2680±0.0483g) were not significantly different to wild type controls (1.3440±0.0297g), being only 6% smaller. In contrast, both *Grb10Δ2-4* KO (1.8140±0.0447g) and *Grb10Δ2-4:Insr* DKO (1.6990±0.0853g) pups were substantially larger than wild type, by 35% (p<0.0001) and 26% (p<0.001), respectively, but not significantly different to each other. Thus, the overgrowth associated with loss of the maternal *Grb10* allele is maintained in DKO pups despite loss of *Insr* expression. A two-way ANOVA test supported this, showing no evidence of an interaction between the genotypes (Figure 5A).

**Figure 5.**
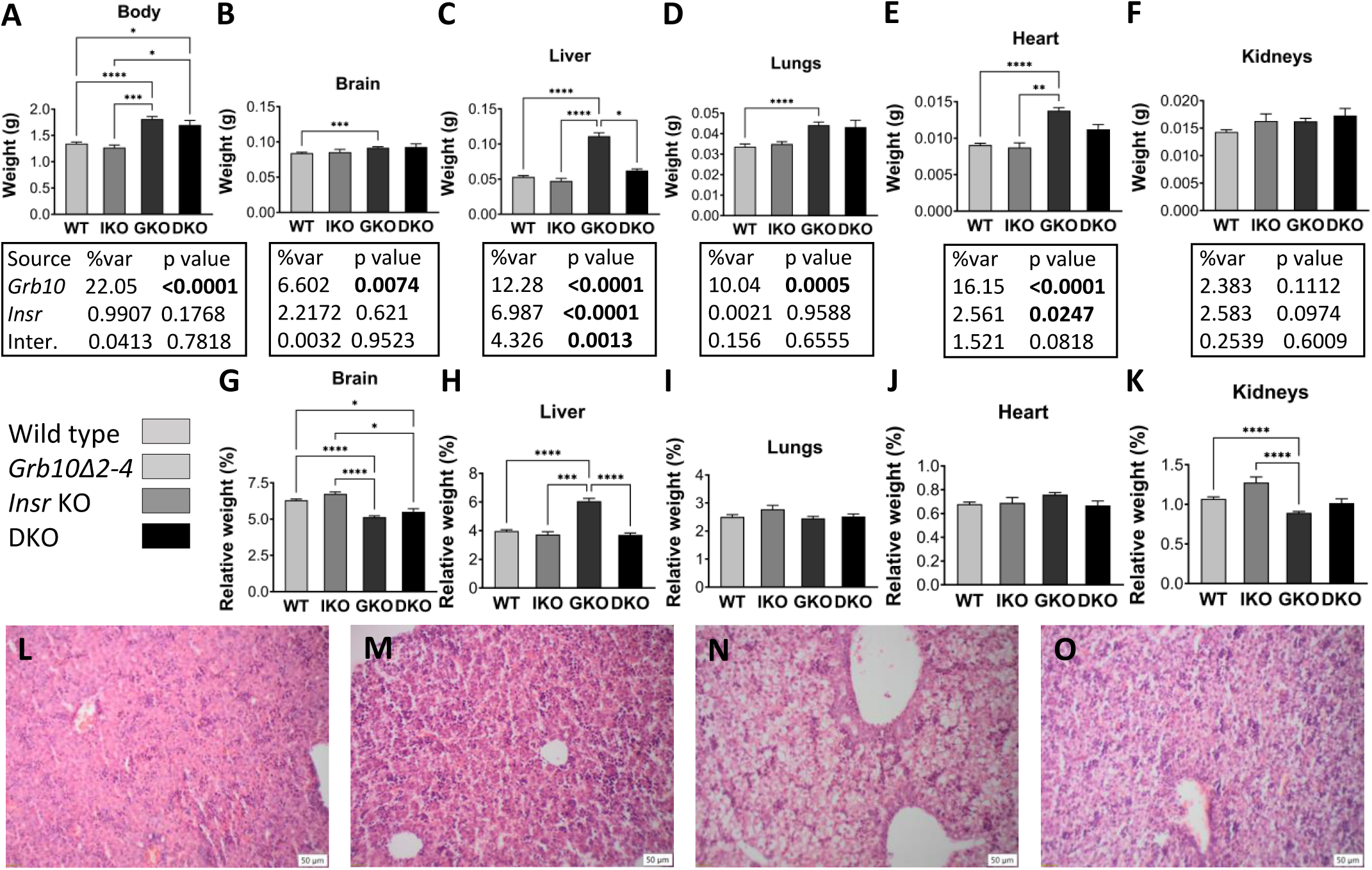
Analyses of whole body and selected dissected organs at PN1 from progeny of crosses between *Grb10Δ2-4* KO and *Insr* KO mice. Data for numerical analyses were pooled into four groups for analysis as described in the Methods, wild type (WT), *Insr* KO (IKO), *Grb10* KO (GKO) and *Grb10*:*Insr* double knockouts (DKO). Body weights are shown for the four offspring genotype groups (A). Actual weights of brain (B), liver (C), lungs (D), heart (E) and kidneys (F) are shown alongside relative weights of the same organs, expressed as a percentage of body mass (G-K). Values represent means and SEM, tested by one-way ANOVA using Kruskal-Wallis and Dunn’s post hoc statistical tests. Summaries of Two-way ANOVA outcomes beneath each graph show the percentage of total variation (%var) and a p value (values significant at p<0.05 in bold) for each source, namely the two single KO genotypes and any interaction (Inter.) between the two. Sample sizes were, WT n=42, *Igf1r* KO n=6, *Grb10* KO n=44, *Grb10*:*Igf1r* DKO n=9. Histological sections of liver, stained with haematoxylin and eosin, are shown for WT (L), *Insr* KO (M), *Grb10* KO (N) and *Grb10*:*Insr* DKO (O) mice. Images are representative of three biological replicates per genotype and were taken at 100x magnification (scale bars show 50μm). Asterisks indicate *p*-values, *p <0.05, **p <0.01, ***p <0.001, ****p<0.0001.

**Table 3.**
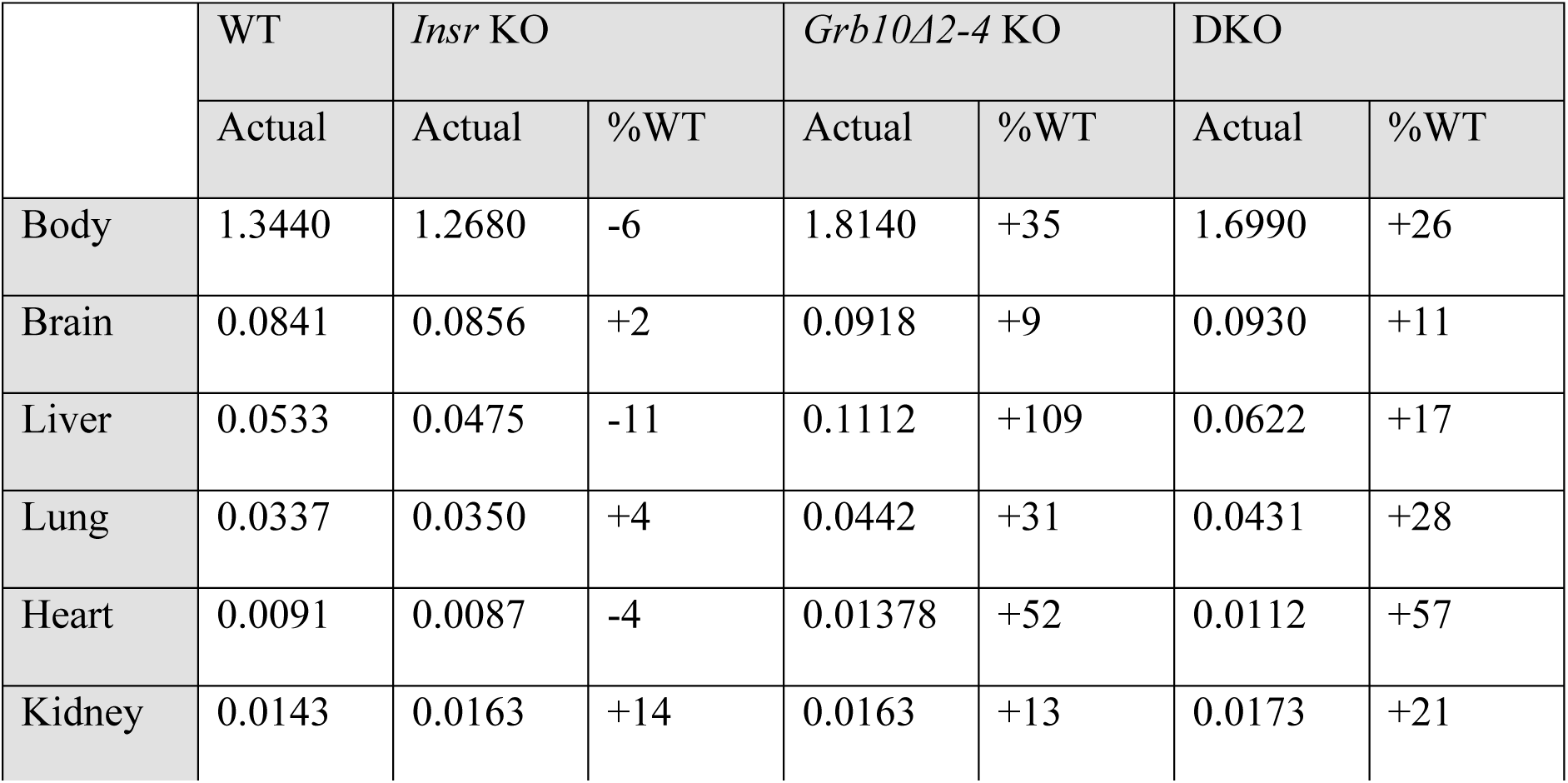
Summary of PN1 body and organ weight data for progeny of crosses between the *Grb10Δ2-4* KO strain and *Insr* KO mice. Mean weights are shown for each genotype together with changes relative to wild type (%WT) for each mutant genotype.

|  | WT | *Insr* KO |  | *Grb10Δ2-4* KO |  | DKO |  |
| --- | --- | --- | --- | --- | --- | --- | --- |
|  | Actual | Actual | % WT | Actual | % WT | Actual | % WT |
| Body | 1.3440 | 1.2680 | -6 | 1.8140 | +35 | 1.6990 | +26 |
| Brain | 0.0841 | 0.0856 | +2 | 0.0918 | +9 | 0.0930 | +11 |
| Liver | 0.0533 | 0.0475 | -11 | 0.1112 | +109 | 0.0622 | +17 |
| Lung | 0.0337 | 0.0350 | +4 | 0.0442 | +31 | 0.0431 | +28 |
| Heart | 0.0091 | 0.0087 | -4 | 0.01378 | +52 | 0.0112 | +57 |
| Kidney | 0.0143 | 0.0163 | +14 | 0.0163 | +13 | 0.0173 | +21 |

#### *Grb10Δ2-4* KO x *Insr* KO offspring PN1 organ mass

As for earlier crosses, the same selection of organs was collected and weighed at PN1 to evaluate body proportions. Organ weights were analysed directly (Figure 5B-F) and as a percentage of total body weight (Figure 5G-K). The patterns of weight differences displayed across the genotypes was consistent with the *Grb10Δ2-4:Insr* DKO pups having an additive phenotype compared with the two single KOs (summarised in Table 3). The brain from *Grb10Δ2-4* KO (0.0918±0.0014g) pups was once again largely spared from the general overgrowth phenotype indicated by body mass, being only 9% larger than wild type (0.0841±0.0014g), which was a significant difference (p<0.01) in this cross (Figure 5B). Brains from *Grb10Δ2-4:Insr* DKO (0.0930±0.0044g) pups were similarly some 11% larger than wild type, whereas *Insr* KO brains (0.0856±0.0039g) were almost indistinguishable at only 2% larger. This meant that *Grb10Δ2-4* KO and *Grb10Δ2-4:Insr* DKO brains were disproportionately small within larger bodies (Figure 5G), compared with wild type (p<0.0001 and p<0.01, respectively) and *Insr* KO (p<0.0001 and p<0.001) brains. Thus, *Grb10Δ2-4:Insr* DKO brain size followed the pattern of the *Grb10Δ2-4* KO and not the *Insr* KO single mutant phenotype.

Liver displayed a particularly interesting pattern of weight differences (Figure 5C). Wild type (0.0533±0.0017g) and *Insr* KO (0.0475±0.0036g) liver sizes were very similar, while *Grb10Δ2-4* KO (0.1112±0.0049g) liver was more than twice normal size, at 109% larger than wild type (p<0.0001), as seen in the previous crosses. However, in this case *Grb10Δ2- 4:Insr* DKO liver (0.0622±0.0023g) was only 17% larger than wild type, and was significantly different (p<0.0001) to *Grb10Δ2-4* KO liver size, but not wild type or *Insr* KO liver, indicating that the disproportionate liver overgrowth associated with loss of *Grb10* expression was largely Insr-dependent. This conclusion was reinforced by the finding that only *Grb10Δ2-4* KO liver was disproportionately enlarged, in comparison with wild type (p<0.0001), *Insr* KO (p<0.0001) and *Grb10Δ2-4:Insr* DKO (p<0.0001). Further, a two-way ANOVA test found an interaction between the genotypes for liver weight (p=0.0013) but not for any other organ (Figure 5A-F). To investigate the liver phenotype further we carried out histological analysis and found that the accumulation of excess lipid previously observed in neonatal *Grb10Δ2-4* KO pups [38] was abrogated in *Grb10Δ2- 4:Insr* DKO pups (Figure 5L-O), indicating that the disproportionate hepatic overgrowth was due to Insr signalling-dependent lipid deposition.

Lungs and heart followed a pattern of size differences like that of body mass. *Grb10Δ2-4* KO (0.0442±0.0015g) and *Grb10Δ2-4:Insr* DKO (0.0431±0.0034g) lungs were similar in size, being 31% (p<0.0001) and 28% (p<0.05) larger, respectively than wild type (0.0337±0.0013g), whereas *Insr* KO (0.0350±0.0011g) lungs were only 4% larger (Figure 5D). Lungs from animals of all four genotypes remained proportionate with body weight (Figure 5I). Similarly, *Grb10Δ2-4* KO (0.01378±0.0004g) and *Grb10Δ2-4:Insr* DKO (0.0112±0.0006g) hearts were both larger than wild type (0.0091±0.0002g) hearts by 52% (p<0.0001) and 24% (p<0.05), respectively, while *Insr* KO (0.0087±0.0006g) hearts were 4% smaller and indistinguishable from wild type (Figure 5E). Hearts from animals of all four genotypes were proportionate with body weight (Figure 5J). In this cross, *Grb10Δ2-4* KO (0.0163±0.0005g) kidneys were 14% (p<0.05) larger than wild type (0.0143±0.0004g) (Figure 5F) but remained disproportionately small (p<0.0001) (Figure 5K). Conversely, *Insr* KO (0.0163±0.0013g) kidneys were 14% larger than wild type and disproportionately large (p<0.01). *Grb10Δ2-4:Insr* DKO (0.0173±0.0013g) kidneys were 21% larger than wild type controls and roughly proportionate such that relative to body mass they were intermediate between the two single KOs. This once again reinforced the sparing of kidneys from the general overgrowth associated with loss of the maternal *Grb10* allele.

#### *Grb10Δ2-4* KO x *Insr* KO offspring e17.5 embryo and placenta

We next investigated the potential for interaction between *Insr* and *Grb10* within the placenta by analysing weights of the whole embryo and placenta at e17.5 (Figure 6). Similar to pups at PN1, compared to wild types (0.9245±0.0240g), *Insr* KO (0.8034±0.0569g) embryos were 13% smaller, though not significantly so, whereas *Grb10Δ2-4* KO embryos (1.3010±0.0445g) and *Grb10Δ2-4:Insr* DKO embryos (1.2130±0.0741g) were larger, by 41% (p<0.0001) and 31%, respectively (Figure 6A). This meant *Grb10Δ2-4* KO (p<0.0001) and *Grb10Δ2-4:Insr* DKO (p<0.05) embryos were both significantly larger than *Insr* KO embryos but not different from each other.

**Figure 6.**
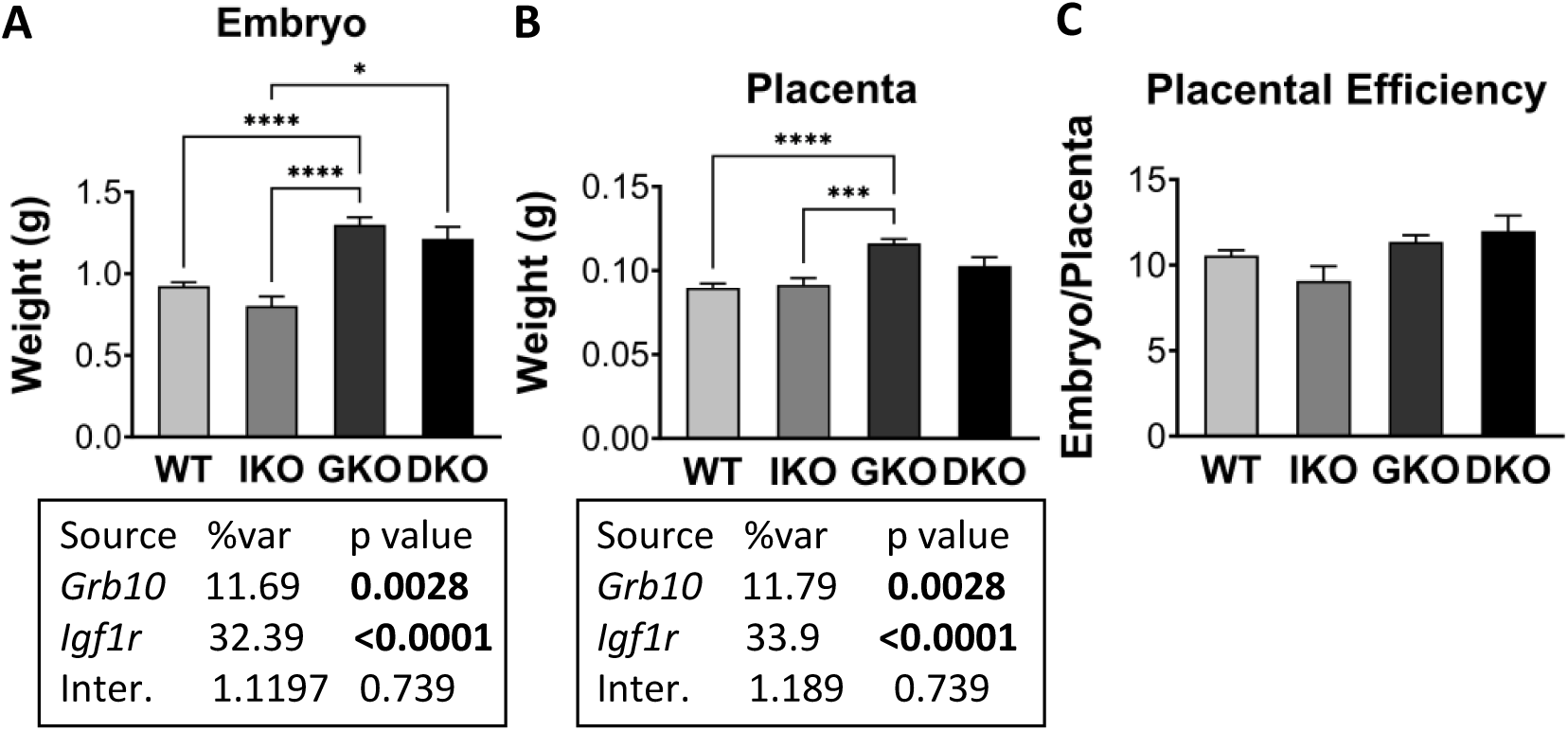
Weight analysis of e17.5 conceptuses from crosses between *Grb10Δ2-4* KO and *Insr* KO mice. Data were pooled into four groups for analysis as described in the Methods, wild type (WT), *Insr* KO (IKO), *Grb10* KO (GKO) and *Grb10*:*Insr* double knockouts (DKO). Weights are shown for the four offspring genotype groups for embryo (A) and placenta (B) and these have been used to calculate the embryo:placenta weight ratio as a measure of placental efficiency (C). Values represent means and SEM, tested by one-way ANOVA using Kruskal-Wallis and Dunn’s post hoc statistical tests. Summaries of Two- way ANOVA outcomes beneath each graph show the percentage of total variation (%var) and a p value (values significant at p<0.05 in bold) for each source, namely the two single KO genotypes and any interaction (Inter.) between the two. Sample sizes were, for WT n=51, *Insr* KO n=13, *Grb10* KO n=52, *Grb10*:*Insr* DKO n=8. Asterisks indicate *p*-values, *p <0.05, ***p <0.001, ****p<0.0001.

In the case of placental weights, wild type (0.0899±0.0024g) and *Insr* KO (0.0915±0.0040g) differed by only 2% while *Grb10Δ2-4* KO (0.1162±0.0026g) and DKO (0.1028±0.0051g) were 29% and 14% larger than wild type, respectively (Figure 6B). The only statistically significant size difference was between *Grb10Δ2-4* KO and either wild type (p<0.0001) or *Insr* KO placentae (p<0.001). When the ratio of embryo to placental mass was calculated as an estimate of placental efficiency, there were no significant differences between genotypes (Figure 4C), though *Grb10* KO (11.35) and DKO (12.0) were slightly higher than wild type (10.55), and *Insr* KO (9.09) slightly lower. Two-way ANOVA found no evidence of an interaction between the genotypes for either embryo (Figure 5A) or placenta (Figure 5B) weight.

#### Survival of *Grb10Δ2-4* KO x *Insr* KO progeny at PN1 and e17.5

Data from the *Grb10Δ2-4* KO x *Insr* KO cross was subject to Chi-squared statistical testing. This indicated that offspring genotype ratios were not significantly different from expected Mendelian ratios at either PN1 (n=101) or e17.5 (n=124) (Supp. Table 2), even though pups lacking *Insr* expression are destined to die within a few days post-parturition of diabetic ketoacidosis [17, 54]. To establish if this was also likely to be true for *Grb10Δ2-4*:*Insr* DKO animals we measured blood glucose levels during dissection of pups on PN1 (Figure 7). Mean glucose concentrations were relatively low and indistinguishable between wild type (2.9mM±0.1) and *Grb10* KO (2.8mM±0.2) animals. Mean glucose levels were also indistinguishable between *Insr* KO (9.1mM±2.5) and DKO (6.4mM±1.7) animals and were significantly higher than wild type (p<0.0001 and p<0.01, respectively) and *Grb10* KO (p<0.0001 and p<0.01, respectively) pups, indicating incipient ketoacidosis in both types of animals lacking *Insr* expression.

**Figure 7.**
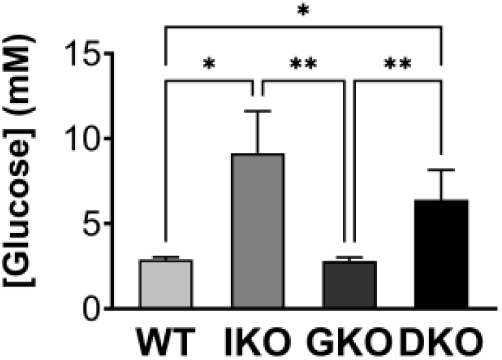
Glucose levels in blood from PN1 progeny from crosses between *Grb10Δ2-4* KO and *Insr* KO mice. Glucose concentration (mM) is shown for progeny of the four genotype groups wild type (WT), *Insr* KO (IKO), *Grb10* KO (GKO) and *Grb10*:*Insr* double knockouts (DKO). Values represent means and SEM, tested by one-way ANOVA using Kruskal-Wallis and Dunn’s post hoc statistical tests. Sample sizes were, for WT n=40, *Insr* KO n=6, *Grb10* KO n=37, *Grb10*:*Insr* DKO n=6. Asterisks indicate *p*-values, *p <0.05, **p <0.01.

## Discussion

Using a mouse genetics approach we have definitively tested whether the Grb10 signalling adaptor protein negatively regulates fetal growth through interaction with either Igf1r or Insr. Growth regulation by Grb10 as a component of the IGF signalling pathway has become the prevailing view because of evidence that Grb10 can physically interact with both receptors [28–30] and can modulate their activity and downstream signalling including *in vivo*, at least in adult mouse tissues [33, 44, 50]. Grb10 activity is modulated through phosphorylation by the raptor kinase, making it a direct target of the mTOR complex [31–33] and further aligning it with Insr signalling. Should an interaction between Grb10 and Insr or Igf1r be responsible for regulation of fetal growth the clear prediction is that mice lacking both Grb10 and either receptor gene will be small at birth to the same extent as the homozygous receptor KO alone, reportedly 60% for *Igf1r* [15] or 90% for *Insr* [17] relative to wild type (Figure 1B). This is because Grb10 will have no influence in the absence of the cognate receptor. However, in crosses between *Grb10* KO and *Igf1r* KO or *Insr* KO mice this was not what we observed and instead the influence of *Grb10* on growth was clearly present in the double knockout offspring both at the level of the whole body, individual organs, and even the gross morphology of *Grb10Δ2-4*:*Igf1r* DKO neonates. This conclusion was strongly supported by two-way ANOVA analysis of body and organ weight data. Across the five datasets presented, evidence of an interaction between the genotypes for body or organ weight was found in only two cases. First, there was evidence of a weak interaction between *Grb10Δ2-4* and *Igf1r* for PN1 body weight (p=0.0135), that was out of line with the four other datasets including that involving the same cross analysed at e17.5. The other exception was more interesting, for PN1 liver weight among offspring of the *Grb10Δ2-4* KO x *Insr* KO, supporting an interaction that explains the disproportionate weight of *Grb10Δ2-4* KO liver through Insr-dependent lipid storage.

In crosses involving the *Igf1r* KO strain and either the *Grb10ins7* or *Grb10Δ2-4* KO strains, the birth weight of DKO pups was closer to that of wild type than either the small *Igf1r* KO or large *Grb10* KO pups. Also, the rate of perinatal lethality and cannibalisation of these DKO mice was much reduced in comparison with that for *Igf1r* KO pups, perhaps reflecting the attainment of an overall size sufficient for a critical function, such as temperature regulation, or the functional rescue of one or more vital organs. Previously, evidence was presented indicating that failure of the *Igf1r* KO lungs to inflate caused death by asphyxia [16] and in support of this we found the *Igf1r* KO lungs to be disproportionately small at 80% lighter than wild type, whereas *Grb10*:*Igf1r* DKO lungs were only some 20-30% smaller than wild type and roughly proportionate with body size. The heart followed a similar pattern, being proportionately reduced in weight in *Igf1r* KO pups, disproportionately enlarged in *Grb10* KO pups and proportionate again in DKO pups, though still heavier than wild type. The dopa decarboxylase gene (*Ddc*), neighbouring *Grb10*, also has a role in promoting growth of the developing heart and is expressed in the developing myocardium, specifically, using a paternally expressed transcript, *Ddc_exon1a* [58]. Despite sharing the imprinting control regions within *Grb10* [59, 60], *Ddc_exon1* and *Grb10* may be expressed in distinct cell populations through the use of separate tissue-specific enhancers [58]. Thus, while the dosage of the two genes is coordinated through genomic imprinting it is not clear whether they regulate fetal heart growth through a shared molecular mechanism.

The brain in *Grb10* KO offspring was essentially spared the general overgrowth of the body, exhibiting relatively small size differences compared to wild type that were inconsistent across the three PN1 datasets examined. Brain weights of *Grb10* KO progeny from the *Grb10ins7* KO x *Igf1r* KO cross were 4% larger than wild type, but not significantly so, 9% (p<0.001) smaller in *Grb10Δ2-4:Insr* KO x *Igf1r* KO cross and 9% (p<0.001) larger in *Grb10Δ2-4* KO x *Insr* KO progeny. Despite this variability, *Grb10* KO brains were always disproportionately small relative to the enlarged body (p<0.0001 in each case). In contrast, the brains of both *Igf1r* KO and *Grb10*:*Igf1r* DKO pups were reduced in size to a similar extent, making clear that changes in brain growth were predominantly due to loss of *Igf1r* expression. This is in keeping with very limited expression of the maternal *Grb10* allele in the developing CNS [36, 37]. This lack of *Grb10* expression means the result could be considered uninformative. However, the paternal *Grb10* allele is strongly expressed in the developing central nervous system and its knockout also has no significant effect on PN1 brain size [36–38], indirectly supporting the idea that Grb10 does not interact with Igf1r to limit fetal brain growth. Kidneys also exhibited sparing in *Grb10* KO offspring across all three crosses, being significantly smaller in relation to body size in all cases. Kidneys in *Igf1r* KO and *Grb10*:*Igf1r* DKO pups were again similarly reduced in size indicating that changes in growth were predominantly due to loss of *Igf1r* expression. However, unlike in brain, maternal *Grb10* is widely expressed in the developing kidney where it is strongest in the epithelial component as judged at the level of both mRNA and protein [37]. In support of a predominantly epithelial role in kidney, human *GRB10* has been shown to be a tumour suppressor in clear cell renal cell carcinoma, a prevalent epithelial kidney cancer [61]. Since kidney growth is driven by expansion of the metanephric mesoderm to fuel nephrogenesis [62], it is possible that in this tissue Grb10 expression is insufficient, that it lacks a cognate growth-mediating receptor, or is functionally redundant with one of the closely-related family members, Grb7 or Grb14 [28]. Grb7 [63] and Grb10 [37], at least, are known to be expressed in the developing kidney and all three adaptor proteins are capable of binding overlapping sets of RTKs [28, 29], but the potential for Grb7 and Grb14 to regulate growth *in vivo* has yet to be explored. Sparing of brain and kidney has been seen in previous crosses involving the *Grb10Δ2-4* KO [37, 38] and *Grb10ins7* [36] strains.

Liver followed an interesting pattern of growth changes across the three crosses. In progeny of both crosses between the *Grb10* KO and *Igf1r* KO strains, *Igf1r* KO liver was reduced in size, albeit to a slightly lesser extent than the body. In contrast, *Grb10* KO livers were disproportionately enlarged, as previously observed [36–38], and *Grb10*:*Igf1r* DKO livers were disproportionately enlarged, to a similar extent. Thus, loss of *Grb10* expression dominated the DKO phenotype, confirming that Grb10 regulates fetal liver size independently of Igfr1. The *Grb10Δ2-4* KO x *Insr* KO cross provided further information. While *Insr* KO offspring had livers of normal size and *Grb10Δ2-4* KO livers were again disproportionately enlarged, those of *Grb10Δ2-4:Insr* DKO offspring were indistinguishable in size from wild type and *Insr* KO livers. Liver histology further revealed that excess lipid accumulation in *Grb10Δ2-4* KO hepatocytes was not seen in *Grb10Δ2-4:Insr* DKO liver, which was indistinguishable from wild type and *Insr* KO livers on this basis. This indicates that during gestation Grb10 normally acts on the Insr to suppresses hepatic lipid storage, perhaps to maximise availability of energy for growth. The result demonstrates for the first time a physiological interaction between Grb10 and the Insr other than in adult tissues [44, 45, 50, 51] . An increase in cell number, mediated by a different tyrosine kinase receptor, as in other tissues, cannot be excluded in the *Grb10* KO liver but is potentially masked by the Insr-mediated hypertrophic expansion of hepatocytes.

Hepatic *Grb10* expression is gradually lost over the first 2-3 weeks after birth and with it the excess weight and lipid accumulation in *Grb10* KO liver [38]. Differentiated adipocytes capable of lipid storage emerge relatively late in development, either in late fetal development (subcutaneous WAT) or in the early post-natal period (gonadal WAT) [64]. Interscapular brown adipose tissue is in place at birth and is important for non- shivering thermogenesis. The transition to energy storage in WAT and utilisation in BAT during the early post-natal period perhaps obviates the need for *Grb10* to suppress hepatic lipid storage and fits with the idea that imprinted genes are important for the transition from maternal dependence to independence [65]. Curiously, in different models of hepatic steatosis *Grb10* expression is induced, including through exposure to cadmium during gestational development [66] or post-natal exposure to tunicamycin or a high fat diet [67]. A liver-specific *Grb10* KO model was used to prove this expression was necessary for steatosis to occur [67]. This indicates a switch in the role of *Grb10* from inhibiting to facilitating hepatic lipid accumulation between fetal and adult life. Using tunicamycin to induce ER stress-mediated steatosis, Luo *et al*., 2018[67] showed that loss of *Grb10* had little effect on insulin-stimulated AKT phosphorylation but significantly down regulated levels of proteins involved in fatty acid synthesis, suggesting involvement of a non- canonical insulin signalling mechanism. Such a pathway has been identified involving insulin- and PI3-kinase-dependent activation of the Snail1 transcription factor in liver [68]. Snail1 binds the fatty acid synthase (FAS) gene promoter, recruiting histone deacetylases and thereby shutting down FAS expression and preventing steatosis. Induction of *Grb10* might therefore be necessary to inhibit this repressive pathway and facilitate lipid accumulation. The absence of Snail1 expression in fetal liver [69] and its presence in adult liver [68] could then explain how loss of *Grb10* expression promotes neonatal steatosis and blocks steatosis in the adult, though this remains to be tested. Steatosis can begin in the fetal or neonatal liver [70] and is recognised as an early indicator of non-alcoholic fatty liver disease, the most prevalent liver disease worldwide [71]. Given the evidence from mouse studies, involvement of *GRB10* in steatosis and NAFLD merits further investigation.

In both crosses involving *Grb10Δ2-4* KO and each of the receptors we evaluated embryo and placental weights at a single late gestational time-point, e17.5, when placental size is maximal. Compared to wild type, we have previously shown that *Grb10Δ2-4* KO conceptuses had a significant difference in mass, evident in the fetus from e12.5 and in the placenta from e14.5 [37]. Also, in a study of wild type litters, *Grb10* expression was found to be higher in the smallest placentae, relative to the largest [72]. Overgrowth of the *Grb10* KO placenta was found to be disproportionate, with greater expansion of the labyrinthine exchange tissue relative to the marginal and junctional zones [57]. This was associated with increased placental efficiency, such that more fetal mass was supported per gram of placental tissue by the *Grb10* KO, likely due to the expanded labyrinthine zone allowing increased nutrient transfer from mother to offspring. Previous studies have concluded that there is no significant difference to wild type in the mass of placentae from *Igf1r* KO, *Insr* KO or even *Igf1r*:*Insr* DKO conceptuses [15, 17]. Our data are consistent with this, although among progeny of the crosses between the *Grb10Δ2-4* KO and *Igf1r* KO strains only a single *Igf1r* KO conceptus was represented, the embryo was approximately 50% smaller than wild type and the placenta only 17% smaller. In both receptor crosses, the mean weights of *Grb10Δ2-4* KO embryos and placentae were significantly larger than wild type, as expected [36–38] and the DKO embryos were intermediate in size. For *Grb10Δ2- 4*:*Igf1r* DKO embryos this meant the embryos were significantly smaller than *Grb10Δ2-4* KO and, although 21% lighter, were not significantly different from wild type. Crucially, *Grb10Δ2-4*:*Igf1r* DKO embryos were not as small as the typical mean mass of *Igf1r* KO embryos reported elsewhere [15, 17], even if only represented by a singleton in our study. The *Grb10Δ2-4*:*Igf1r* DKO placentae were also intermediate in size, between *Grb10Δ2-4* KO and wild type, but without being significantly different to either. While there was a significant difference in placental efficiency only between *Grb10Δ2-4* KO and *Grb10Δ2- 4*:*Igf1r* DKO conceptuses, there was a non-significant trend towards increased *Grb10Δ2-4* KO placental efficiency in both crosses examined, as in our earlier report [57]. This result alone is ambiguous but the balance of evidence against involvement of Igf1r in controlling placental size [15, 17] favours the interpretation that Grb10 controls growth independently of Igf1r in the placenta as well as the embryo.

*Insr* KO progeny from our crosses did not display a significant growth deficit, in terms of whole-body mass at PN1 (-6%) or e17.5 (-13%), or in the mass of any individual PN1 organs. This at first appears to contrast with a reported 10% growth deficiency in e18.5 *Insr* KO progeny of an *Insr* KO x *Igf1r* KO cross [17], where the numbers of embryos weighed (n = 121, including 9 *Insr^-/-^*) were very similar to our PN1 sample size (n = 101, including 9 *Insr^-/-^*). However, it should be noted that the relatively high significance level (p<0.05) was obtained using a student’s *t* test without any correction for multiple testing. That said, the fact that *Insr* KO pups were consistently smaller by 6-13% across 3 different crosses and two separate studies, suggests the impact of *Insr* KO on fetal growth could be biologically relevant. Indeed, it seems feasible to assume that disruption in energy regulation should impact fetal growth and perhaps surprising that such an effect is not more obvious. In agreement with a previous report [17], *Insr* KO had no impact on placental weight. The lack of a clear growth deficit associated with *Insr* KO did not affect the interpretation of our data since the well characterised overgrowth of *Grb10* KO pups was still evident in *Grb10Δ2-4*:*Insr* DKO pups, ruling out Insr as a major receptor through which Grb10 mediates fetal growth regulation. This was evident through examination of individual organ weights as well as whole body weights. Most straightforwardly, lungs and heart were clearly enlarged to a similar extent in *Grb10Δ2-4* KO and *Grb10Δ2-4*:*Insr* DKO pups and differed significantly from both wild type and *Insr* KO organs. In this cross, *Grb10Δ2-4* KO brain and kidneys again exhibited sparing from the general overgrowth of the body, which meant there were only small weight differences across the genotypes for these organs, though both *Grb10Δ2-4* KO and *Grb10Δ2-4*:*Insr* DKO brain and kidneys were disproportionately small relative to the whole-body overgrowth exhibited by pups of these genotypes. At PN1 there was no obvious deficit in the number of *Insr* KO or *Grb10Δ2-4*:*Insr* DKO pups but both had significantly elevated blood glucose levels, indicative of incipient ketoacidosis, as previously observed for *Insr* KO neonates [54]. In summary, the *Grb10Δ2-4* KO x *Insr* KO cross data has established that increased fetal growth associated with loss of maternal *Grb10* expression is not mediated through interaction with the *Insr*. Any impact of the *Insr* alone on fetal growth regulation is modest and instead it is primarily or solely a regulator of glucose homeostasis, which we have shown extends to facilitating lipid storage in the fetal liver, which is normally inhibited by *Grb10*. The role of insulin in the control of hepatic glucose regulation is well documented [73].

As well as optimising body size during fetal growth, the growth of individual tissues and organs must be coordinated to achieve a size compatible with efficient function. Tissue proportions can be influenced by the environment. For instance, when nutrient supply is limited during development proportions can be altered in order to preserve brain growth over other organs in animals ranging from *Drosophila* to human (see [74]) which has been termed brain sparing. By limiting growth in only peripheral tissues *Grb10* could, therefore, be an important determinant of brain sparing. More generally, our work shows how body proportions, as well as size, is altered through the actions of two independent growth regulatory pathways. Although we have not identified the ‘growth’ receptor, or receptors, on which Grb10 acts, the findings allow us to make some important inferences. In at least two pathways growth and energy homeostasis are intimately linked through Insr and mTOR signalling. While it was initially anticipated that *Grb10* would prove to be the third imprinted gene influencing the Ins/IGF signalling pathway, we have shown instead that imprinting has evolved to influence more than one growth regulatory pathway. Theories for the evolution of imprinting, including the conflict hypothesis, tend to focus on individual genes rather than pathways. It is generally agreed that the benefits of voluntarily shutting down one of the two parental alleles must outweigh the cost, most obviously the risk of losing the one active copy but also, once adapted to the single gene dose, the risk of the silent copy becoming active. These risks may explain why more genes are not subject to imprinting within a single pathway, with the consequences of the resulting imbalances amply illustrated by imprinting disorders such as BWS and SRS [12].

Our data highlight that the coordination of organ size regulation during fetal development can be disrupted through maternal *Grb10* KO in a manner that is not apparent through disruption of *Igf1r* expression. In *Igf1r* KO PN1 pups, organs derived primarily from each of the three germs layers, ectoderm (brain), mesoderm (heart, kidneys) and endoderm (liver, lungs) were all reduced in size. This is consistent with Igf1r, which mediates signalling of Igf1 and Igf2 [15, 16], impacting growth during early embryogenesis. A study of *Igf2* KO embryos supports this, finding that disruption of cell proliferation and survival in a narrow window between e9-e10, resulted in significant changes in cell number, detectable from e11 [75], which is a few days earlier than a difference in mass can be properly discerned [15, 37, 75]. This window coincides with the early post gastrulation period when there is rapid expansion of the three germ layers and the initial events in organogenesis are taking place. We predict that by acting within a similar developmental window and engaging with one or more different receptors, Grb10 influences growth of a more limited set of tissue lineages. One possibility is that Grb10 acts on lineage-specific progenitors as they emerge during early organogenesis, since their expansion is known to regulate organ size as demonstrated, for instance, by genetic ablation experiments (e.g. [76]). Further work will be needed to identify the receptor(s) with which Grb10 interacts to influence fetal growth. Candidates must be expressed in a pattern overlapping with *Grb10* during embryogenesis, be capable of binding Grb10 and will themselves have a role in fetal growth regulation. Although many RTKs are known to regulate relevant processes, such as proliferation, survival and differentiation of specific cell types, their impact on overall body size and tissue proportions is less well studied. Growth deficiency phenotypes of RTK knockout mice can be challenging to investigate, for instance, where there are pleiotropic effects such as patterning defects, gestational lethality or potentially because of redundancy with related family members. Fundamental understanding of fetal growth regulation has potential benefits for the development of novel interventions that improve neonatal outcomes and life-long health for the wider population, including those with rare growth disorders.

## Declarations

### Ethical approvals

Experiments involving mice were subject to local ethical review by the University of Bath Animal Welfare and Ethics Review Board and carried out under licence from the United Kingdom Home Office. The manuscript has been written as closely as possible in accordance with the Animal Research: Reporting of In Vivo Experiments (ARRIVE) guidelines (https://arriveguidelines.org/).

### Consent for publication

Not applicable.

### Availability of data and materials

The datasets used and analysed during the current study are available from the corresponding author on reasonable request.

### Author contributions

AW conceived the project, collected and analysed data, wrote the manuscript and assembled the figures. FMS and ASG set up genetic crosses, collected data and contributed to data analysis. KM performed histological examination of the liver, carried out the statistical analyses and generated the final graphs and images. All authors read and approved the final manuscript.

## Acknowledgements

For generously supplying mouse strains we thank Domenico Accili (*Insr* KO) and Martin Holzenberger (*Igf1r* KO). We are grateful to University of Bath Biological Services Unit staff for outstanding animal care.

---

**Supplementary Table 1.**
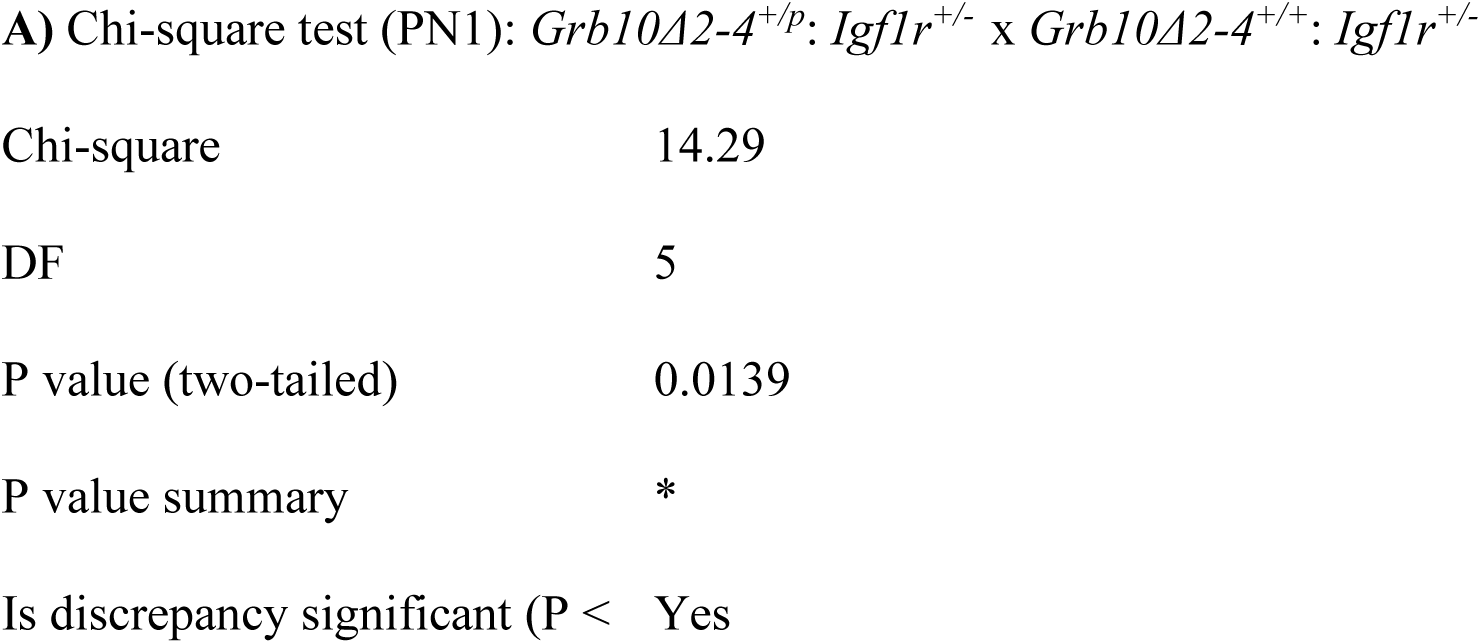
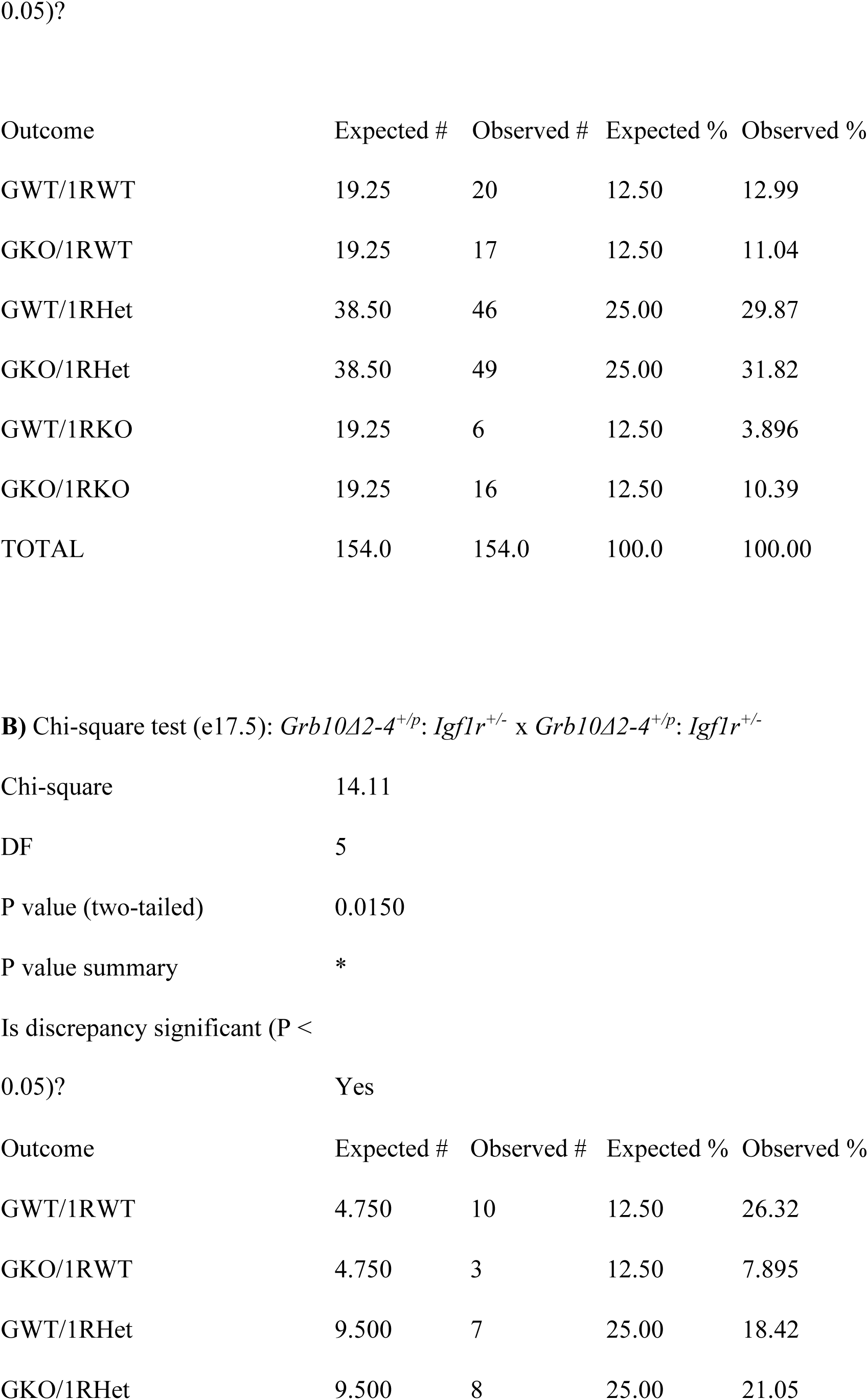
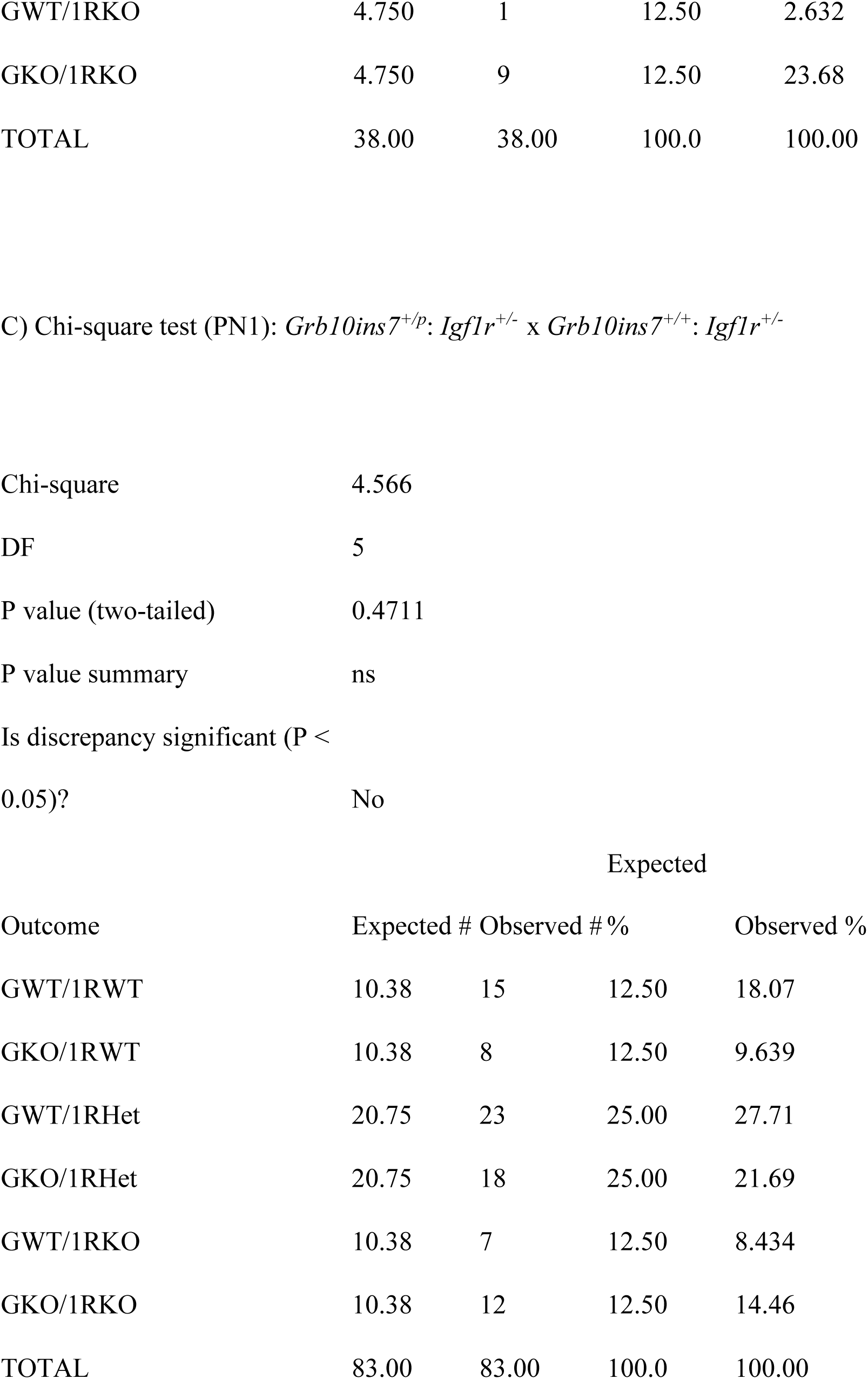
Chi-squared statistical tests of offspring survival from crosses involving *Grb10* KO and *Igf1r* KO strains. Offspring collected from crosses between *Grb10Δ2-4^+/p^*: *Igf1^+/-^* females and *Grb10Δ2-4^+/+^*: *Igf1^+/-^* males at, (A) PN1 and (B) e17.5. (C) Offspring collected at PN1 from crosses between *Grb10ins7^+/+^*: *Igf1^+/-^* females and *Grb10ins7^+/p^*: *Igf1^+/-^* males. Deviation from the expected Mendelian ratio was considered significant at p<0.05.

**Supplementary Table 2.**
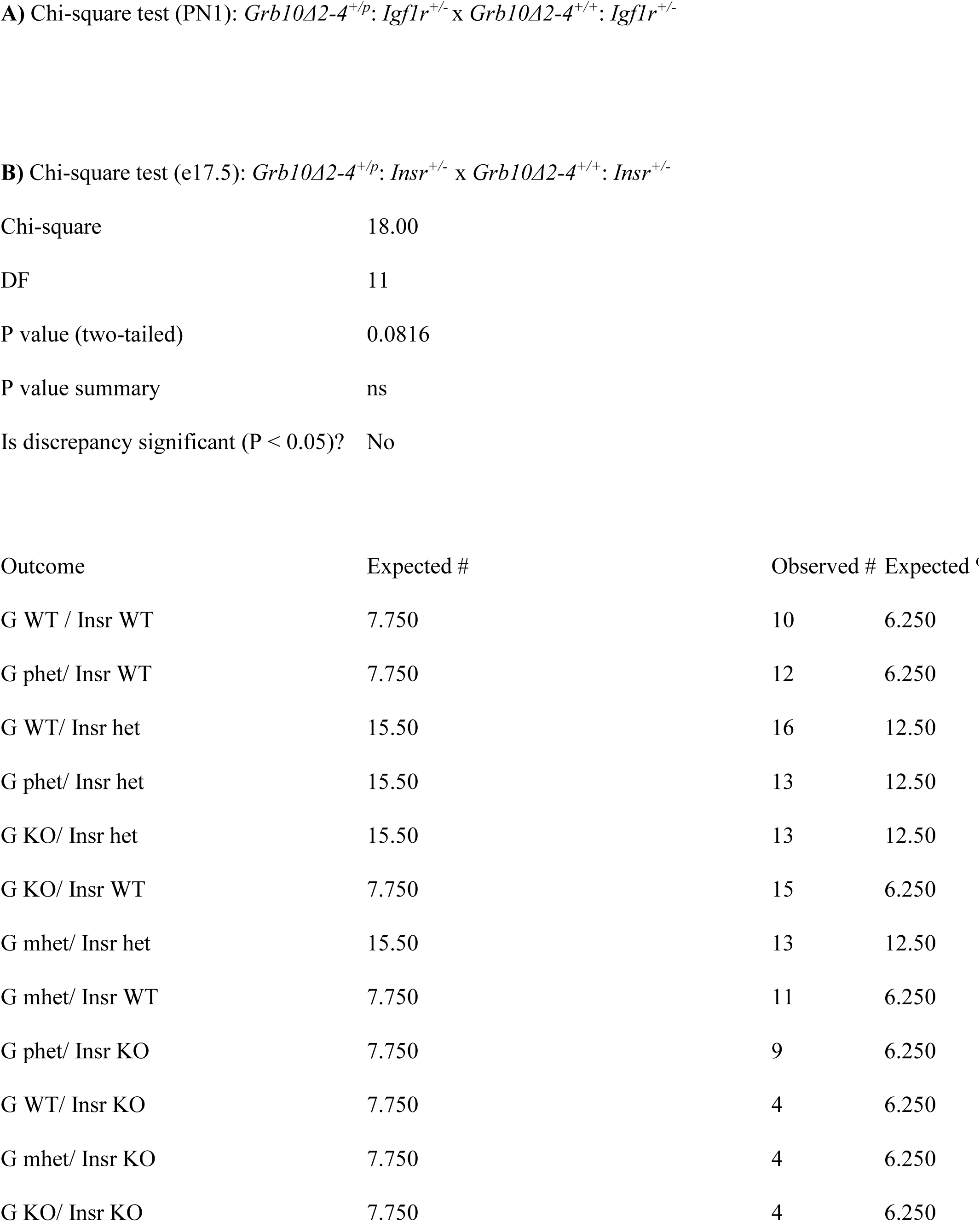
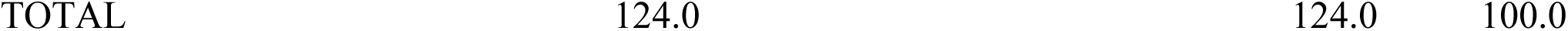
Chi-squared statistical tests of offspring survival from crosses between the *Grb10 Δ2-4* KO and *Insr* KO strains. Offspring collected from crosses between *Grb10Δ2-4^+/p^*: *Insr^+/-^* females and *Grb10Δ2-4^+/p^*: *Insr^+/-^* males at PN1 (A) and at e17.5 (B). Deviation from the expected Mendelian ratio was considered significant at p<0.05.

