## Supplemental Tables for "Imprinted *Grb10*, encoding growth factor receptor bound protein 10, regulates fetal growth independently of the insulin-like growth factor type 1 receptor (*Igf1r*) and insulin receptor (*Insr*) genes"

**Supplementary Tables**

**A)** Chi-square test (PN1): *Grb10Δ2-4*$^{+/p}$: *Igf1r*$^{+/-}$ x *Grb10Δ2-4*$^{+/+}$: *Igf1r*$^{+/-}$

| | |
|---|---|
| Chi-square | 14.29 |
| DF | 5 |
| P value (two-tailed) | 0.0139 |
| P value summary | * |
| Is discrepancy significant (P < 0.05)? | Yes |

| Outcome | Expected # | Observed # | Expected % | Observed % |
|---|---|---|---|---|
| GWT/1RWT | 19.25 | 20 | 12.50 | 12.99 |
| GKO/1RWT | 19.25 | 17 | 12.50 | 11.04 |
| GWT/1RHet | 38.50 | 46 | 25.00 | 29.87 |
| GKO/1RHet | 38.50 | 49 | 25.00 | 31.82 |
| GWT/1RKO | 19.25 | 6 | 12.50 | 3.896 |
| GKO/1RKO | 19.25 | 16 | 12.50 | 10.39 |
| TOTAL | 154.0 | 154.0 | 100.0 | 100.00 |

**B)** Chi-square test (e17.5): *Grb10Δ2-4*$^{+/p}$: *Igf1r*$^{+/-}$ x *Grb10Δ2-4*$^{+/p}$: *Igf1r*$^{+/-}$

| | |
|---|---|
| Chi-square | 14.11 |
| DF | 5 |
| P value (two-tailed) | 0.0150 |
| P value summary | * |
| Is discrepancy significant (P < 0.05)? | Yes |

| Outcome | Expected # | Observed # | Expected % | Observed % |
|---|---|---|---|---|
| GWT/1RWT | 4.750 | 10 | 12.50 | 26.32 |
| GKO/1RWT | 4.750 | 3 | 12.50 | 7.895 |
| GWT/1RHet | 9.500 | 7 | 25.00 | 18.42 |
| GKO/1RHet | 9.500 | 8 | 25.00 | 21.05 |
| GWT/1RKO | 4.750 | 1 | 12.50 | 2.632 |
| GKO/1RKO | 4.750 | 9 | 12.50 | 23.68 |
| TOTAL | 38.00 | 38.00 | 100.0 | 100.00 |

C) Chi-square test (PN1): *Grb10ins7*$^{+/p}$: *Igf1r*$^{+/-}$ x *Grb10ins7*$^{+/+}$: *Igf1r*$^{+/-}$

| | |
|---|---|
| Chi-square | 4.566 |
| DF | 5 |

| | | | | |
|---|---|---|---|---|
| P value (two-tailed) | 0.4711 | | | |
| P value summary | ns | | | |
| Is discrepancy significant (P < 0.05)? | No | | | |
| Outcome | Expected # | Observed # | Expected % | Observed % |
| GWT/1RWT | 10.38 | 15 | 12.50 | 18.07 |
| GKO/1RWT | 10.38 | 8 | 12.50 | 9.639 |
| GWT/1RHet | 20.75 | 23 | 25.00 | 27.71 |
| GKO/1RHet | 20.75 | 18 | 25.00 | 21.69 |
| GWT/1RKO | 10.38 | 7 | 12.50 | 8.434 |
| GKO/1RKO | 10.38 | 12 | 12.50 | 14.46 |
| TOTAL | 83.00 | 83.00 | 100.0 | 100.00 |

**Supplementary Table 1.** Chi-squared statistical tests of offspring survival from crosses involving *Grb10* KO and *Igf1r* KO strains. Offspring collected from crosses between *Grb10Δ2-4*$^{+/p}$: *Igf1*$^{+/-}$ females and *Grb10Δ2-4*$^{+/+}$: *Igf1*$^{+/-}$ males at, (A) PN1 and (B) e17.5. (C) Offspring collected at PN1 from crosses between *Grb10ins7*$^{+/+}$: *Igf1*$^{+/-}$ females and *Grb10ins7*$^{+/p}$: *Igf1*$^{+/-}$ males. Deviation from the expected Mendelian ratio was considered significant at p<0.05.

**A)** Chi-square test (PN1): *Grb10Δ2-4*$^{+/p}$: *Igf1r*$^{+/-}$ x *Grb10Δ2-4*$^{+/+}$: *Igf1r*$^{+/-}$

**B)** Chi-square test (e17.5): *Grb10Δ2-4*$^{+/p}$: *Insr*$^{+/-}$ x *Grb10Δ2-4*$^{+/+}$: *Insr*$^{+/-}$

| | |
|---|---|
| Chi-square | 18.00 |
| DF | 11 |
| P value (two-tailed) | 0.0816 |
| P value summary | ns |
| Is discrepancy significant (P < 0.05)? | No |

| Outcome | Expected # | Observed # | Expected % |
|---|---|---|---|
| G WT / Insr WT | 7.750 | 10 | 6.250 |
| G phet/ Insr WT | 7.750 | 12 | 6.250 |
| G WT/ Insr het | 15.50 | 16 | 12.50 |
| G phet/ Insr het | 15.50 | 13 | 12.50 |
| G KO/ Insr het | 15.50 | 13 | 12.50 |
| G KO/ Insr WT | 7.750 | 15 | 6.250 |
| G mhet/ Insr het | 15.50 | 13 | 12.50 |
| G mhet/ Insr WT | 7.750 | 11 | 6.250 |
| G phet/ Insr KO | 7.750 | 9 | 6.250 |
| G WT/ Insr KO | 7.750 | 4 | 6.250 |
| G mhet/ Insr KO | 7.750 | 4 | 6.250 |
| G KO/ Insr KO | 7.750 | 4 | 6.250 |
| TOTAL | 124.0 | 124.0 | 100.0 |

**Supplementary Table 2.** Chi-squared statistical tests of offspring survival from crosses between the *Grb10 Δ2-4* KO and *Insr* KO strains. Offspring collected from crosses between *Grb10Δ2-4*$^{+/p}$: *Insr*$^{+/-}$ females and *Grb10Δ2-4*$^{+/p}$: *Insr*$^{+/-}$ males at PN1 (A) and at e17.5 (B). Deviation from the expected Mendelian ratio was considered significant at $p<0.05$.
